# Experimental Time Points Guided Transcriptomic Velocity Inference

**DOI:** 10.64898/2026.02.17.705910

**Authors:** Xinyuan Zang, Xin Shu, Nina Zhang, Yu Wu, Mingyue Deng, Xiaolong Zhou, Jie Yang, Chen-Yu Zhang, Xiaoyong Wang, Zhen Zhou, Jin Wang

## Abstract

Time-series single-cell RNA sequencing enables longitudinal tracking of biological processes, yet cellular trajectory reconstruction informed by experimental time remains challenging. Existing trajectory inference methods either perform *de novo* reconstruction without leveraging experimental time points, or prioritize transitions between time points while paying less attention to intra-time-point dynamics. To reconcile experimental time points with local precision, we present CellDyc, a semi-supervised learning framework that leverages experimental time-point supervision to reconstruct transcriptomic velocities and recover an intrinsic gene-embedded time. CellDyc consistently outperforms existing approaches in reconstructing cellular trajectories across development, disease, and reprogramming contexts. Biologically, CellDyc provides novel insights, such as resolving temporal heterogeneity in erythroid maturation and quantitatively demonstrating that the immunosuppressive environment delays monocyte differentiation in glioblastoma. CellDyc integrates seamlessly with downstream tools like CellRank and remains robust even when only inferred temporal information is available. Collectively, CellDyc offers a rigorous, data-driven solution for deciphering time-resolved cellular dynamics.

## Introduction

The advent of time-series single-cell RNA sequencing (scRNA-seq) has provided a powerful capability to track critical biological processes, including development, disease progression, and cellular reprogramming. Time-series designs overcome the limited temporal coverage of single-time-point sampling by systematically collecting samples at multiple key experimental time points, thus enabling longitudinal tracking [1-11]. However, due to the destructive nature of scRNA-seq, the resulting data remains a series of discrete, static snapshots. Trajectory inference methods have thus emerged to resolve the continuous biological processes underlying these snapshots. By bridging the gap between discrete sampling and continuous cellular dynamics, these methods significantly deepen our understanding of cell fate decisions and their underlying molecular mechanisms [12-19]. Despite these advances, existing methods still fall short of effectively leveraging experimental time points for the precise reconstruction of cellular dynamics.

One major category of methods, including pseudotime [20-25] and RNA velocity [26-30], typically disregards the explicit information provided by experimental time points. These tools pool cells from all time points to reconstruct the temporal dimension and trajectory direction *de novo*: they order cells along a trajectory based on transcriptomic similarity and prior biological knowledge, or estimate transcriptomic velocity (the time derivative of gene expression states) via RNA splicing kinetic signals. While valuable for providing high-resolution cellular dynamics, these methods are often sensitive to data noise and parameter selection [31, 32]. Furthermore, their heavy reliance on prior assumptions can compromise their reliability across diverse biological contexts.

In contrast, Optimal Transport (OT)-based methods [33-40] explicitly leverage the chronological order of experimental time points, thereby ensuring highly reliable global directionality. However, these approaches generally treat each time point as a static probability distribution of cell states. By focusing solely on transitions between time points while ignoring cellular dynamics within them, OT-based methods are often confined to describing macroscopic developmental trends, failing to reconstruct high-resolution, instantaneous cellular dynamics [41].

To bridge this gap, we propose CellDyc, a semi-supervised learning framework that utilizes experimental time-point labels to predict transcriptomic velocity directly from transcriptomic profiles. Unlike previous approaches, CellDyc integrates explicit temporal information from experimental time points with the complete heterogeneity of cell states, both within and across time points. By translating these inputs into temporal relationships and transcriptomic differences between a cell and its nearest neighbors, CellDyc achieves accurate inference of instantaneous single-cell dynamics via transcriptomic velocity prediction. Furthermore, proceeding from the premise that gene expression evolves over time—and conversely, that “real-world” time is embedded within gene expression patterns—we developed a Gene Clock module within CellDyc. This module uses experimental time points as external supervision to recover gene-embedded time, an biologically grounded, intrinsic coordinate system that provides a precise temporal reference for biological processes.

We validated CellDyc across diverse biological contexts spanning development, disease, and reprogramming. These include embryogenesis in *Caenorhabditis elegans (C. elegans)* [8] and zebrafish [9], glioblastoma-associated monocyte differentiation [42], erythrocyte maturation during gastrulation [10], and mouse embryonic fibroblast reprogramming [11]. In all benchmarks, CellDyc consistently outperformed both OT-based methods (Waddington-OT [39], Moscot [40]) and RNA velocity approaches (scVelo [27], cellDancer [28]). Notably, CellDyc yielded novel biological insights, revealing that the immunosuppressive microenvironment of glioblastoma delays monocyte differentiation and recovering temporal heterogeneity among cell lineages during erythrocyte maturation.

Finally, CellDyc is designed for seamless integration with existing computational infrastructure. Its predicted transcriptomic velocity vectors are compatible with downstream analysis tools originally developed for RNA velocity methods. For instance, in our reprogramming study, CellDyc vectors were directly integrated into CellRank [43, 44] to calculate cell fate probabilities, demonstrating the framework’s broad applicability and potential.

## Results

### Inferring Transcriptomic Velocity and Gene-Embedded Time Through Semi-Supervised Learning

CellDyc is a semi-supervised learning framework that leverages experimental time points to predict transcriptomic velocity and encode intrinsic gene-embedded time. Experimental time points offer partial supervision for predicting continuous time, while integrating this temporal information with gene expression similarity supplies partial guidance for inferring transcriptomic velocity. This paradigm of learning from partially labeled data is naturally addressed through semi-supervised approaches. We decompose the problem into two coupled subtasks: (1) constructing a continuous cellular timeline and (2) inferring instantaneous directional trends, each adapted with dedicated semi-supervised strategies (Fig. 1b).

**Figure 1.**
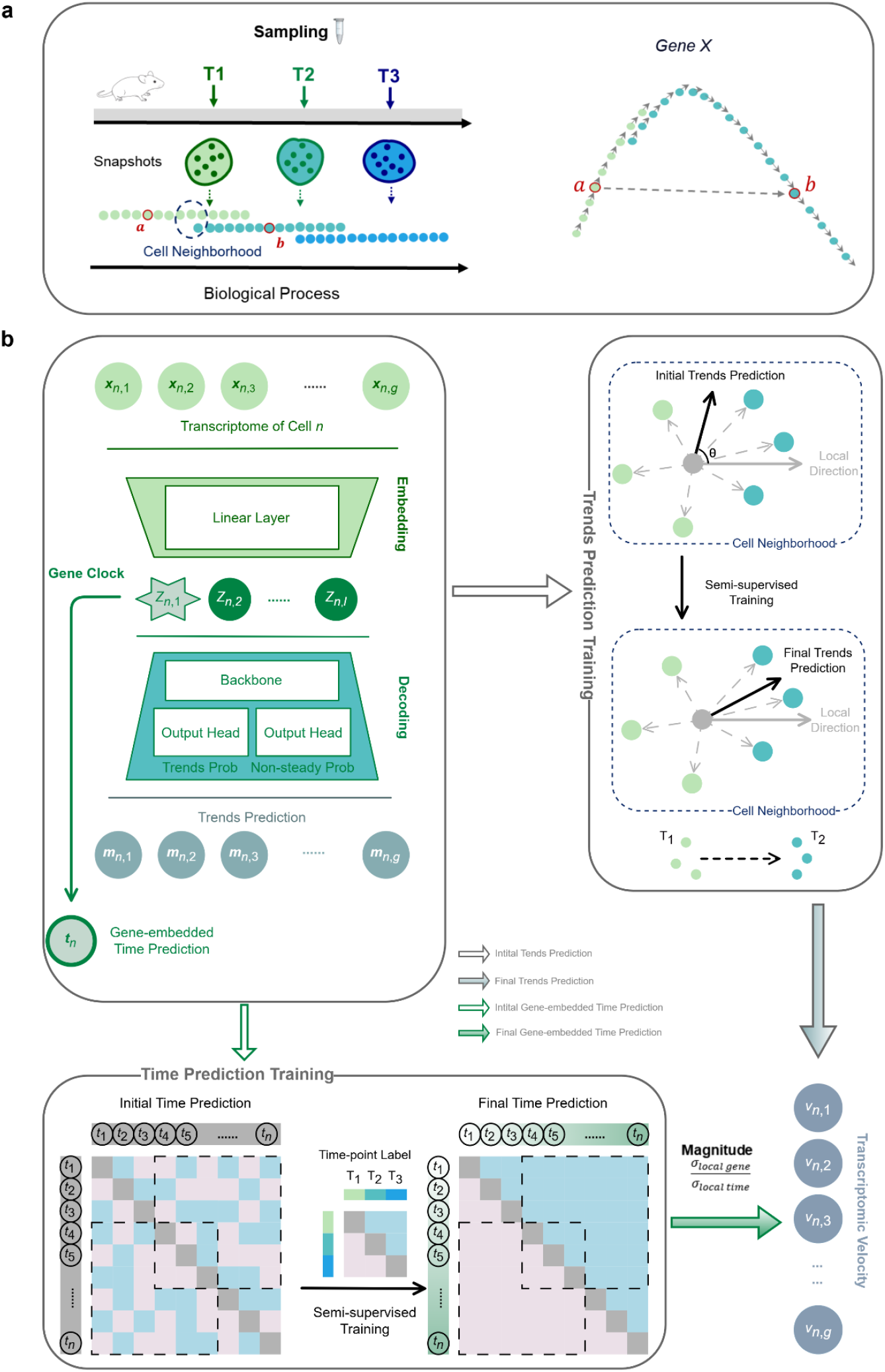
Overview of the CellDyc framework. **a**, Schematic representation of time-series single-cell RNA sequencing data. Left: Samples collected at multiple key experimental time points (T1, T2, T3) capture a series of static snapshots of cell states within a biological process, where cell states from adjacent time points often overlap in state space. Right: Cell state transitions between time points do not necessarily reflect the true instantaneous cellular dynamics. **b**, The CellDyc architecture and training strategy. Left (Model Architecture): The unified framework takes single-cell transcriptomes (*x*_*n*_) as input. A linear embedding layer encodes a latent Gene Clock (*z*_*n*,1_) alongside other features. The decoder predicts a Directional Trends Probability Vector and a Non-steady State Probability Vector to generate the trends prediction (*m*_*n*_). The Gene Clock is specifically used for gene-embedded time prediction (*t*_*n*_) and modulates the magnitude of the final velocity. Top Right (Trends Prediction Training): Illustration of the semi-supervised trends prediction task. For a given cell (grey), “local directionality” is computed by weighting directed difference vectors to its neighbors (blue and orange cells) based on their temporal orientation (T1 vs. T2). This integrates experimental time information with local transcriptomic similarity to supervise the trends prediction. Bottom (Time Prediction Training): Illustration of the time prediction task. The model learns to reconstruct a continuous timeline from discrete time-point labels (T1, T2, T3). The matrices show pairwise rankings of cells, where blue indicates the row cell precedes the column cell, and red indicates the reverse. Through semi-supervised training, the “Initial Gene-embedded time” (disordered) is refined into a “Final Gene-embedded time” that preserves temporal rank constraints, generating a continuous ordering. Bottom Right: The final transcriptomic velocity prediction combines the magnitude (derived from the ratio of local gene variance to local time variance) with the predicted trends.

While experimental time points capture long-term transcriptional evolution, transcriptomic velocity reflects instantaneous cellular changes. Existing methods such as Waddington-OT [39] and Moscot [40] directly map cell fates between discrete time points—an approach limited by the inherently complex and nonlinear nature of biological processes. As illustrated in Fig. 1a, cell state transitions inferred merely between time points do not necessarily reflect true instantaneous dynamics; the difference between short-term and long-term dynamics is not merely one of magnitude, but fundamentally one of direction within gene expression space.

To resolve this, we integrate experimental time with local gene expression similarity by focusing on cell-specific neighborhoods (a focal cell and its nearest neighbors). Directed difference vectors between a central cell and its neighbors are weighted by temporal orientation (e.g., T1 vs. T2) to derive “local directionality,” which serves as the primary supervisory signal for the trend prediction task (Fig. 1b, Top Right). Since neighborhoods exhibit varying temporal spans—with some containing isochronous cells that preclude direction inference—we implement adaptive weighting schemes that enable robust learning from incomplete directional labels (Methods).

Though experimental time points provide only coarse-grained supervision, pairwise comparisons generate rank relationships that prove sufficiently informative for ordering most cell pairs. We learn a linear combination of genes—the Gene Clock—that maximally preserves these temporal rank constraints. This module reconstructs continuous gene-embedded time from discrete time-point labels, trained to achieve a final temporal order that respects the ranking matrix (Fig. 1b, Bottom). The linear formulation of the Gene Clock deliberately constrains model capacity, thereby mitigating overfitting to labeled rank constraints while preserving generalizability to unlabeled cell pairs.

Both tasks are unified within a single architecture featuring a linear embedding layer whose first dimension constitutes the Gene Clock. This clock operates concurrently in two capacities: (1) as an independent representation of gene-embedded time with a dedicated loss function, and (2) as part of the latent vector for directional trend prediction. Furthermore, the Gene Clock calibrates the magnitude of the final transcriptomic velocity by calculating the ratio of local gene variance to local time variance; this magnitude is combined with the directional trend prediction to yield the transcriptomic velocity prediction (Fig. 1b, Bottom Right).

### CellDyc Robustly Recovers Transcriptomic Velocity and Time Across Varying Sampling Densities

We utilized scDesign3 to generate simulated datasets with ground-truth transcriptomic velocities and time to benchmark CellDyc’s performance [45]. While scDesign3 does not directly output velocity vectors, it models gene expression as a time-dependent function via Generalized Additive Models for Location, Scale, and Shape (GAMLSS). We therefore derived ground-truth transcriptomic velocities by applying finite difference methods to these fitted models (Methods). We first constructed a continuous reference dataset comprising 5,000 cells and 500 genes (Fig. 2a), and then simulated a time-series experiment by sampling cells at designated experimental time points according to a normal distribution (Fig. 2b, Methods).

**Figure 2.**
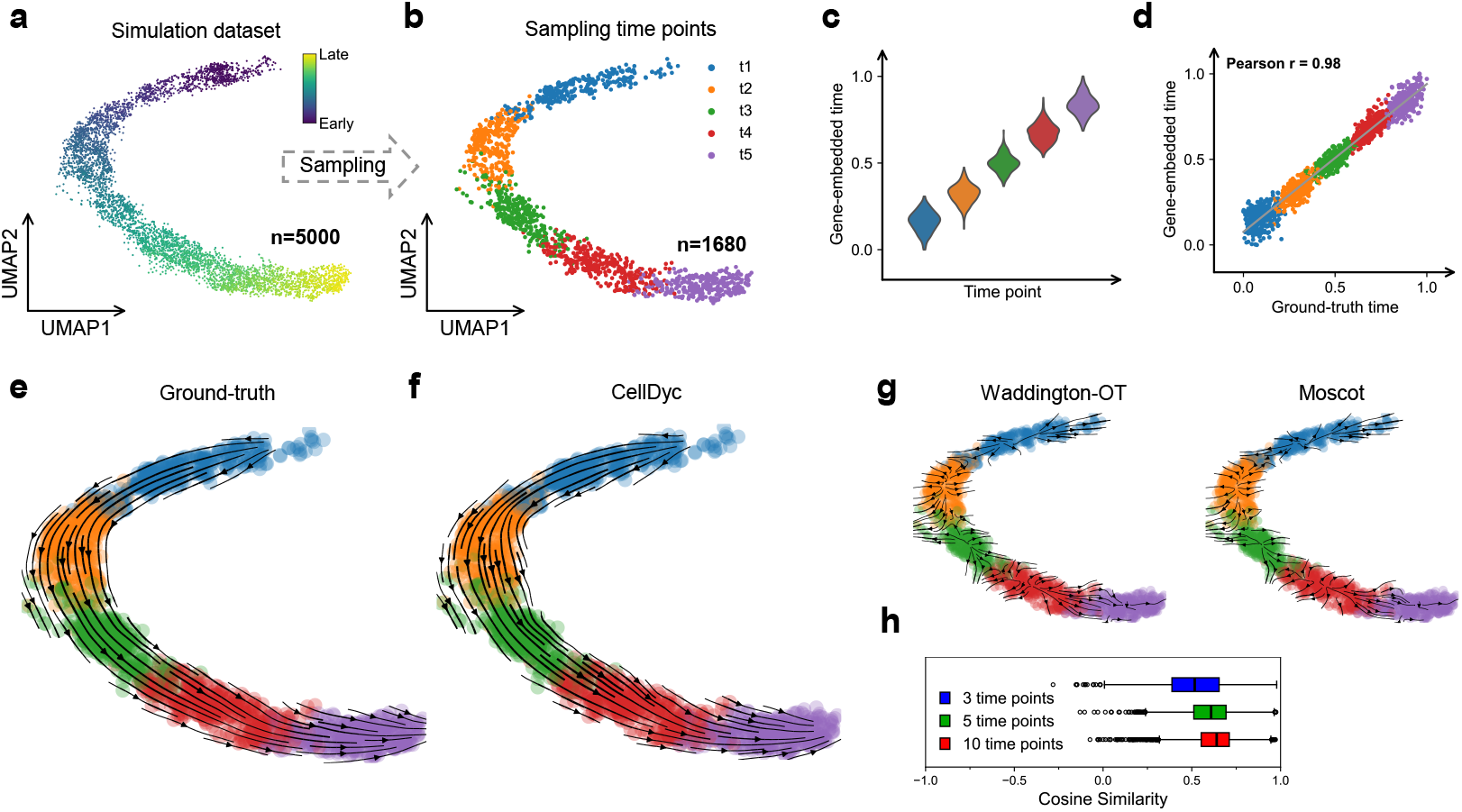
Validation of CellDyc in recovering ground-truth time and transcriptomic velocity using simulated data. **a**, Simulation dataset comprising 5,000 cells. **b**, Temporal sampling at five time points according to a normal distribution, resulting in 1,680 cells. **c**, Violin plots of predicted gene-embedded time across sampling time points. **d**, The correlation between predicted gene-embedded time and the ground-truth time, Pearson’s correlation coefficient is indicated. **e,f**, Projection of transcriptomic velocities onto UMAP. (**e**) Ground-truth velocities. (**f**) CellDyc-predicted velocities. **g**, Projection of cell state transitions onto UMAP: (left) Waddington-OT, (right) Moscot. **h**, Cosine similarity between each cell’s ground-truth and CellDyc-predicted velocities across varying temporal sampling densities. Box plots display medians (vertical lines) and interquartile ranges (boxes) for three sampling conditions.

Evaluation using a five-time-point design revealed that this sampling density effectively captured the continuous cell state landscape. CellDyc’s Gene Clock module inferred gene-embedded time that exhibited precise correspondence with the input experimental time points (Fig. 2c) and showed a high correlation with the hidden ground-truth time (Pearson’s r = 0.98) (Fig. 2d). Visualizing the ground-truth transcriptomic velocities on a Uniform Manifold Approximation and Projection (UMAP) embedding confirmed a clear directional temporal progression (Fig. 2e). Comparative analysis demonstrated that CellDyc accurately predicted instantaneous transcriptomic velocities that closely recapitulated the ground truth (Fig. 2f). In contrast, OT-based methods—Waddington-OT [39] and Moscot [40]—struggled to recover even the general trajectory direction at this sampling density (Fig. 2g), likely because they rely on macroscopic transitions between time points rather than learning fine-grained dynamics within them.

We further assessed robustness across varying sampling densities. CellDyc’s performance remained consistent with a denser sampling of 10 time points (Extended Data Fig. 1b–e). Under this high-resolution condition, Waddington-OT and Moscot showed marked improvement, capturing overall trajectory patterns despite residual inaccuracies (Extended Data Fig. 1f). This suggests their performance depends heavily on minimal gaps between time points to reduce the conflation of long-term and short-term state changes. Conversely, a sparse design with only 3 time points resulted in incomplete coverage of the cell state landscape (Extended Data Fig. 1g). Remarkably, CellDyc maintained robust predictive accuracy under these constrained conditions, displaying only modest performance attenuation and demonstrating strong resilience to limited sampling (Extended Data Fig. 1h–k). In comparison, Waddington-OT and Moscot exhibited further performance degradation (Extended Data Fig. 1l).

To quantify the impact of sampling density, we computed the cosine similarity between the predicted and ground-truth transcriptomic velocity vectors for each cell—a metric specifically applicable to CellDyc, as Waddington-OT and Moscot output probabilistic transition matrices rather than explicit velocity vectors. Although accuracy declined slightly as time points decreased, CellDyc remained reliable even with only 3 sampling points (Fig. 2h). While these simulations preclude universal sampling density recommendations for all biological contexts, they confirm that increased temporal resolution generally enhances performance. Specifically, for this synthetic system, five time points represented an optimal balance between accuracy and experimental efficiency.

### CellDyc Accurately Reconstructs High-Resolution Developmental Dynamics in *C. elegans* and Zebrafish

To benchmark the reliability of CellDyc in real-world biological systems, we applied it to two canonical model organisms: *C. elegans* and zebrafish. Both species offer unique advantages for developmental studies, including optically transparent embryos that enable direct visualization of embryogenesis. Notably, *C. elegans* embryos possess a remarkably small number of invariant cell lineages that have been nearly completely resolved [46].

We first evaluated CellDyc using *C. elegans* embryonic data from Packer et al. [8], who integrated scRNA-seq profiles across multiple experimental time points and mapped them onto the invariant lineage tree. By aligning each cell’s expression profile with a high-resolution whole-embryo RNA-seq time series, the authors estimated a precise “embryo time” for individual cells. Following the data selection strategy of Wang et al. [47], we focused on the AB lineage. Since the dataset pooled cells from numerous embryos, most lineage nodes were represented multiple times. We retained only embryos with unambiguous sampling times and randomly selected a single representative cell per lineage node using a specially designed sampling strategy to preserve the original temporal distribution (Methods). This curation yielded 333 cells spanning 3 experimental time points, each annotated with precise embryo time and fully resolved lineage relationships (Fig. 3a, b).

**Figure 3.**
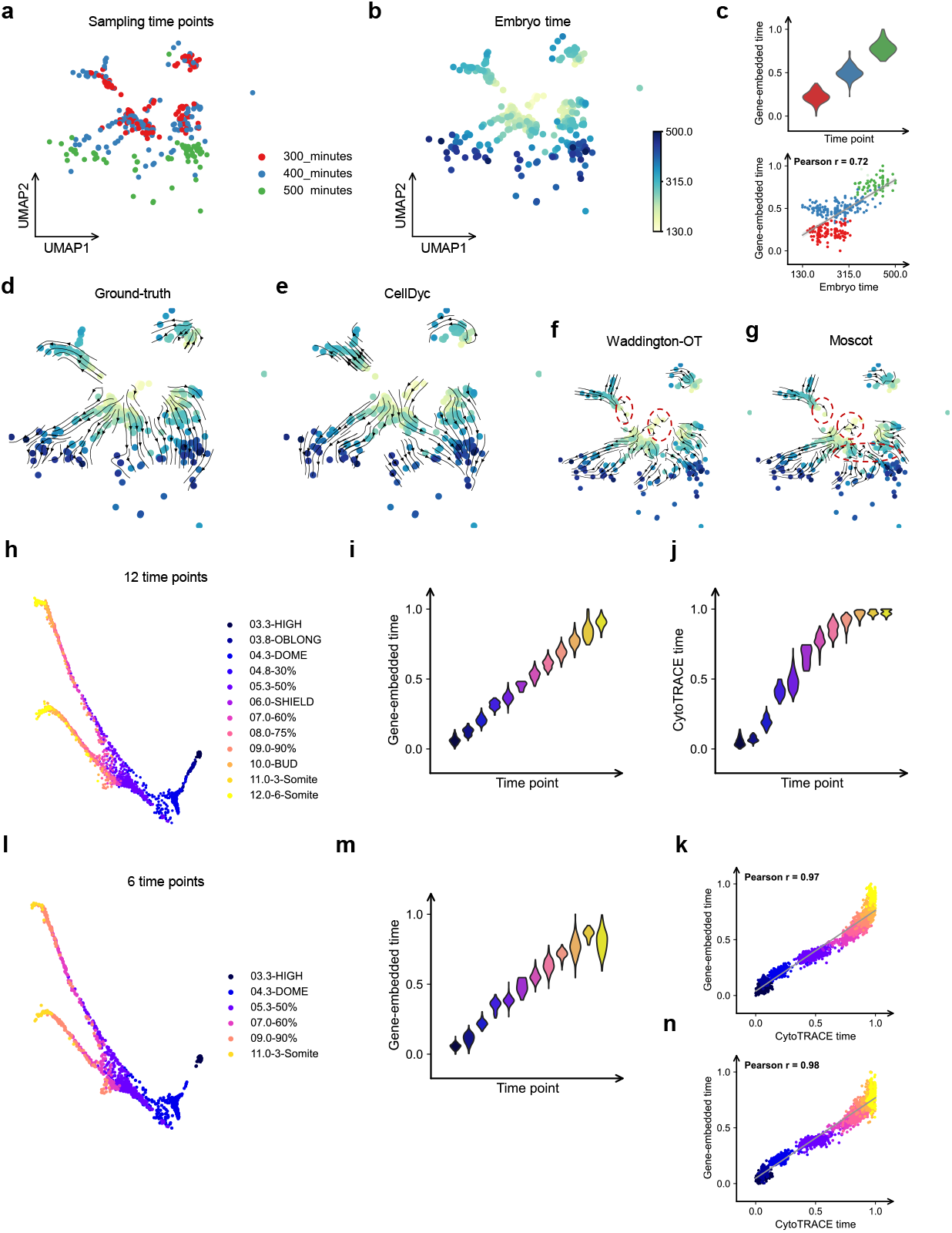
Reconstruction of transcriptomic velocity and masked time points in *C. elegans* and zebrafish embryogenesis. **a,b**, UMAP embedding of 333 cells from the AB lineage of *C. elegans* embryos. Each cell corresponds to a node in the lineage tree, colored by sampling time points (**a**) and embryo time (**b**). **c**, Top panel: Violin plots of predicted gene-embedded time across sampling time points. Bottom panel: Pearson’s correlation coefficients (r) between predicted and embryo time. **d,e**, Projection of transcriptomic velocities onto UMAP. (**d**) Ground-truth velocities. (**e**) CellDyc-predicted velocities. **f,g**, Projection of cell state transitions onto UMAP: (**f**) Waddington-OT, (**g**) Moscot. **h**, Force-directed layout of 2,341 cells of the axial mesoderm lineage during zebrafish embryogenesis, colored by sampling time points. **i**, Violin plots of predicted gene-embedded time across sampling time points from the full 12-time-point model. **j**, Violin plots of CytoTRACE time (1-CytoTRACE score) across sampling time points. **l**, Six time points selected for CellDyc training are shown. **m**, Violin plots of predicted gene-embedded time across sampling time points from the selected 6-time-point model. **k,n**, The correlation between predicted gene-embedded time and the CytoTRACE time, Pearson’s correlation coefficient is indicated (**k**) 12-time-point model; (**n**) 6-time-point model.

Ground-truth transcriptomic velocities were derived by directly comparing expression profiles between parent and daughter cells in the lineage tree. The projection of these ground-truth velocities onto the UMAP embedding faithfully recapitulated the differentiation trajectory along embryo time (Fig. 3d). Remarkably, CellDyc recovered gene-embedded time using only the sparse sampling time points (Fig. 3a). This inferred gene-embedded time not only aligned with the provided sampling times (Fig. 3c, top) but also accurately recovered the high-resolution embryo time that was withheld from the model (Pearson’s r = 0.72; Fig. 3c, bottom). Furthermore, CellDyc’s predicted transcriptomic velocities faithfully reproduced the differentiation trajectory revealed by the ground truth (Fig. 3e). In comparison, Waddington-OT [39] and Moscot [40] successfully captured the core developmental trajectory, despite exhibiting some minor local inaccuracies (Fig. 3f, g). This suggests that the limited number of cells in this *C. elegans* dataset provided a sufficiently fine sampling density that, despite having only three time points, enabled OT-based methods to infer global trends—a finding consistent with observations from high-density sampling in simulated data.

We next assessed CellDyc’s performance on a zebrafish embryogenesis atlas spanning 12 experimental time points (3.3–12 hours post-fertilization) [9]. Specifically, we analyzed the axial mesoderm lineage subset (encompassing notochord and prechordal plate trajectories) extracted by Lange et al. [43], which contains 2,341 cells retaining all 12 original time points (Fig. 3h). We examined the capability of CellDyc to reconstruct unobserved time points using this dataset. As a baseline, CytoTRACE estimates developmental potential based on the number of detectably expressed genes per cell [48]; in developmental contexts, CytoTRACE time (1 – CytoTRACE Score) serves as a proxy for intrinsic cellular time.

When provided with all 12 time points, CellDyc’s gene-embedded time accurately reproduced the sampling time distribution (Fig. 3i) and slightly outperformed CytoTRACE time in discriminating the three latest time points (10–12h) (Fig. 3j). Overall, gene-embedded time showed a strong linear correlation with CytoTRACE time (Fig. 3k). To test the model’s capacity for temporal interpolation, we trained CellDyc on a subset of only 6 time points and evaluated its performance on the full 12-time-point dataset (Fig. 3l). Although temporal discrimination of the 10–12h window decreased slightly (Fig. 3m), the correlation with CytoTRACE time remained robust and even improved marginally (Fig. 3n). A systematic evaluation including 9 and 3 time points revealed that while reduced temporal sampling progressively impaired the discrimination of late developmental stages (Extended Data Fig. 2a, b, d, e), the correlation with CytoTRACE time remained largely stable (Extended Data Fig. 2c, f).

Finally, we examined how temporal sparsity affected CellDyc’s prediction of transcriptomic velocities. With all 12 time points, velocity projections perfectly matched the expected biological trajectories. Reducing the input to 6 time points caused a loss of trajectory resolution between 3–4h, and with only 3 time points, performance declined further, correctly capturing only the local trajectory from 5–10h (Extended Data Fig. 2g). In stark contrast, Waddington-OT and Moscot failed to recover the major trajectory even when provided with the full 12 experimental time points (Extended Data Fig. 2h), underscoring CellDyc’s superior ability to reconstruct high-resolution transcriptomic velocities from discrete experimental snapshots.

### CellDyc Recovers Gene-Embedded Time to Unveil Delayed Differentiation Kinetics in Tumor Microenvironments

We further sought to extend CellDyc to scenarios beyond conventional sampling-time annotations. Zman-seq, an in vivo timestamping technique pioneered by Kirschenbaum et al. [42], sequentially labels circulating immune cells via intravenous injection of fluorophore-conjugated CD45 antibodies. Upon tissue infiltration, cells become shielded from subsequent antibody injections, enabling the inference of tissue residence time from their fluorescent signatures. This approach was employed to dissect immune remodeling dynamics mediated by TREM2 antagonistic antibodies (aTREM2) in glioblastoma models. The dataset captures temporal trajectories of monocyte-to-tumor-associated macrophage (TAM) differentiation under two conditions: control IgG treatment, which recapitulates an immunosuppressive microenvironment driving monocytes toward Arg1^+^Gpnmb^+^ immunosuppressive TAMs, and aTREM2 treatment, which redirects differentiation toward Acp5^+^ pro-inflammatory TAMs [49].

Because Zman-seq labeling is not instantaneous, it introduces additional technical asynchrony on top of the inherent biological asynchrony of cell populations. This results in noisy, highly asynchronous experimental time labels (Fig. 4a), which complicates standard trajectory reconstruction. Nevertheless, by leveraging these labels to predict transcriptomic velocities, CellDyc successfully delineated two distinct monocyte-to-TAM trajectories—Arg1^+^Gpnmb^+^ versus Acp5^+^ TAMs—under the respective treatment conditions (Fig. 4b). In stark contrast, both Waddington-OT [39] and Moscot [40] failed to reconstruct biologically meaningful trajectories from the experimental time data (Fig. 4c). Consistent with this difficulty, Kirschenbaum et al. noted that scVelo did not faithfully reconstruct the expected trajectories on this data either.

**Figure 4.**
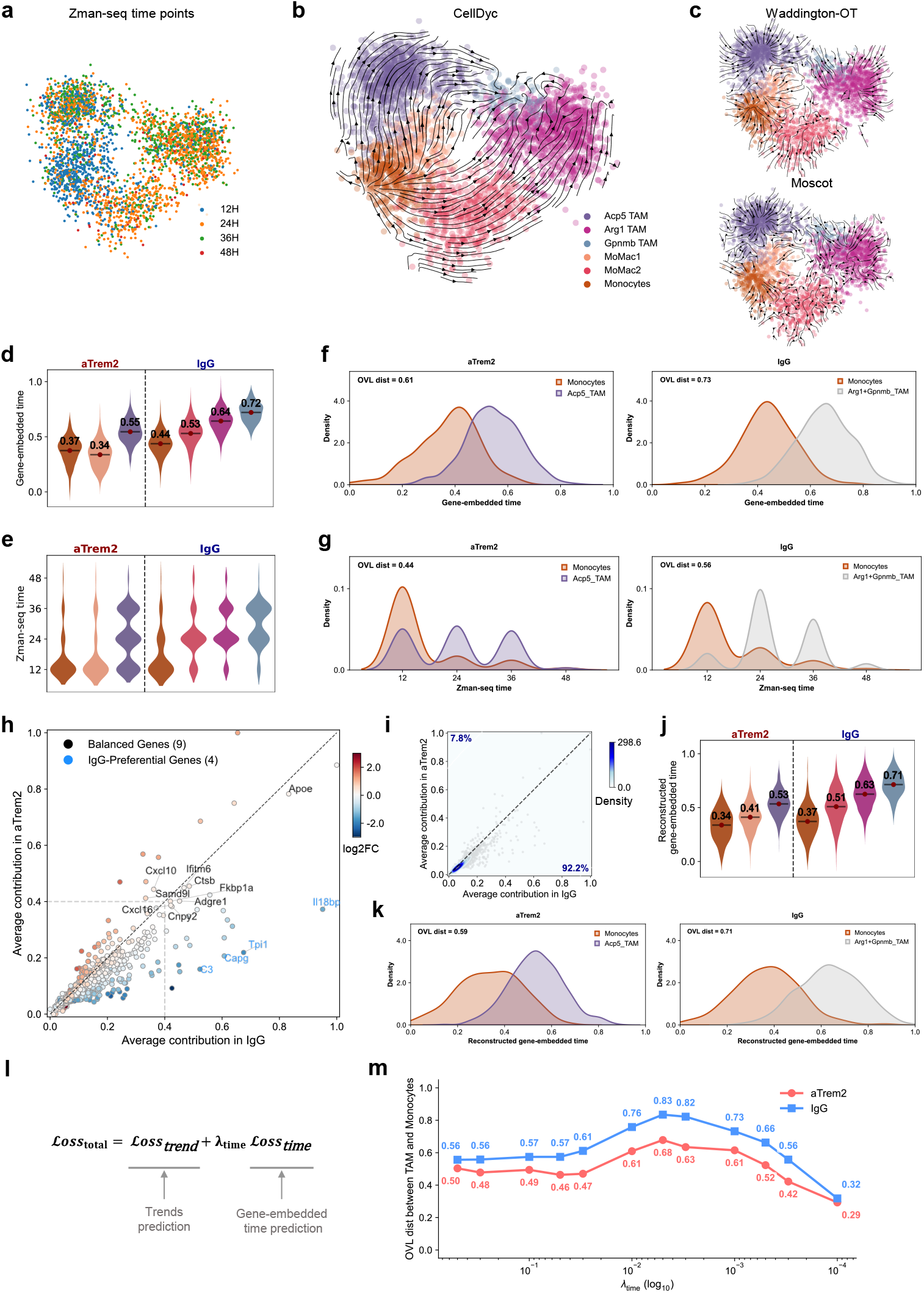
Characterization of monocyte differentiation trajectories and microenvironmental influences using CellDyc. **a**, Metacell graph projection of 3108 cells from monocyte-to-TAM differentiation, colored by time points from Zman-seq. **b**, Transcriptomic velocities derived from CellDyc are projected onto the metacell graph, revealing two distinct monocyte-to-TAM trajectories under the two treatment conditions. **c**, In contrast to (**b**), Waddington-OT (top) and Moscot (bottom) failed to infer biologically meaningful cell trajectories. **d**, Violin plots of predicted gene-embedded time across cell types under aTREM2 treatment versus IgG control conditions. **e**, Violin plots of observed Zman-seq time across cell types under aTREM2 treatment versus IgG control conditions. **f**, Distributions of predicted gene-embedded time in monocytes and TAMs under aTREM2 treatment versus IgG control conditions, with OVL distance between TAMs and monocytes indicated. Left: aTREM2 treatment; Right: IgG control. **g**, Distributions of observed Zman-seq time in monocytes and TAMs under aTREM2 treatment versus IgG control conditions, with OVL distance indicated. Left: aTREM2 treatment; Right: IgG control. **h**, Scatter plot showing the contribution of individual genes to gene-embedded time under two treatment conditions. Each dot represents a single gene. The x-axis denotes each gene’s contribution to gene-embedded time in the IgG-treated group, and the y-axis denotes its contribution to gene-embedded time in the aTREM2-treated group. Among genes with high contributions (>0.4 in either condition), nine “Balanced Contribution Genes” (located near the diagonal, indicating similar contributions in both groups) and four “IgG-Preferential Genes” (located farther from the diagonal toward the x-axis, indicating stronger contribution in the IgG group relative to aTREM2) are highlighted on the plot. **i**, Density plot showing gene contributions to gene-embedded time, similar to (**h**) in terms of x- and y-axis definitions. The percentages of genes located above and below the diagonal are annotated on the plot, indicating that, overall, genes contributing to gene-embedded time tend to show stronger contributions in the IgG-treated group. **j**, Violin plots of gene-embedded time reconstructed using all 13 selected genes (nine “Balanced Contribution Genes” and four “IgG-Preferential Genes”) across cell types under aTREM2 treatment versus IgG control conditions. **k**, Distributions of reconstructed gene-embedded time (as in (**j**)) in monocytes and TAMs under aTREM2 treatment versus IgG control conditions, with OVL distance between TAMs and monocytes indicated. Left: aTREM2 treatment; Right: IgG control. **l**, Schematic illustration of the loss function in CellDyc. The hyperparameter *λ*_*tttt*_ controls the weight of the time-loss component. **m**, Line plot showing the OVL distances between TAMs and monocytes across different *λ*_*tttt*_ settings under aTREM2 treatment compared to IgG control conditions. Each condition is represented by a separate line. The x-axis represents *λ*_*tttt*_ values on a logarithmic scale (log10).

To understand this capability, we compared CellDyc’s recovered gene-embedded time against the original experimental time (observed Zman-seq time). While the experimental time distributions for each cell type spanned all four time points, the gene-embedded time distributions were markedly more focused, revealing sharper temporal separation between cell types (Fig. 4d-e). This indicates that the gene-embedded time learned by CellDyc provides a more informative temporal representation than the raw experimental labels. We quantified the temporal segregation between TAMs and monocytes using the Overlapping Coefficient (OVL; defined as 1-OVL for distance; see Methods). The results demonstrated that gene-embedded time consistently outperformed experimental time in distinguishing cell states across both treatment groups (OVL distances: 0.61 vs. 0.44 for aTREM2; 0.73 vs. 0.56 for IgG) (Fig. 4f-g).

Notably, Arg1^+^ and Gpnmb^+^ TAMs exhibited evidently higher mean gene-embedded time than Acp5^+^ TAMs (Fig. 4d). The increased OVL distance between TAMs and monocytes in the IgG group compared to the aTREM2 group (0.73 vs. 0.61; Fig. 4f) suggests that monocyte differentiation is temporally protracted within the immunosuppressive microenvironment. While calculations based on experimental time displayed a similar trend (Fig. 4g), the inherent asynchrony of the raw labels obscured this relationship visually. This microenvironment-mediated effect on differentiation kinetics represents a largely unexplored dimension, with comparable phenomena only recently documented in cell line systems [49].

We next dissected the mechanisms underlying this temporal encoding by quantifying each gene’s contribution to the gene-embedded time in both treatment groups. The Gene Clock module displayed a marked bias toward the IgG condition, with 92.2% of genes weighted more heavily for reconstructing time under IgG treatment (Fig. 4h-i). From the high-contribution genes, we identified nine “Balanced Contribution Genes” (located near the diagonal) and four “IgG-Preferential Genes” (distal to the diagonal). We then tested whether gene-embedded time could be reconstructed using only these 13 genes and their associated weights. The resulting reconstructed gene-embedded time nearly matched the performance of the full model (OVL distances: 0.59 vs. 0.61 for aTREM2; 0.71 vs. 0.73 for IgG) and similarly captured the delayed differentiation kinetics characteristic of immunosuppressive conditions (Fig. 4j-k).

Further analysis revealed that exclusive encoding with Balanced Contribution Genes effectively distinguished TAMs from monocytes but failed to resolve the IgG-specific differentiation delay (Extended Data Fig. 3a-b). Conversely, IgG-Preferential Genes captured the inter-group temporal differences but could not sufficiently separate TAMs from monocytes within the aTREM2 group (Extended Data Fig. 3c-d). These results demonstrate the utility of mining Gene Clock-derived gene combinations as candidate markers.

Finally, to understand how CellDyc leverages the interaction between its two prediction tasks to move beyond the supervision provided by the time labels, we examined its dual-loss training architecture, where the hyperparameter λ_*time*_ modulates the weight of the time loss (Fig. 4l). Systematic variation of λ_*time*_ revealed that both excessively high and low values impair the establishment of accurate temporal relationships (Fig. 4m). This indicates that gene-embedded time encoding relies on integrating the time prediction task and the trends prediction task, rather than depending solely on either component. The time loss appears to serve as a scaffold, designating a latent space for explicit temporal encoding. Consequently, extreme λ_*time*_ values lead the model to either merely replicate the noisy experimental time (when too large) or produce time-agnostic latent variables (when too small) (Extended Data Fig. 4).

### CellDyc Resolves Temporal Heterogeneity and Establishes a Temporal Reference in Erythroid Maturation

We evaluated CellDyc on a mouse gastrulation erythropoiesis scRNA-seq dataset comprising eight experimental time points that capture the entire erythroid lineage specification, from hematoendothelial progenitors to Erythroid 3 cells (Fig. 5a) [10]. By projecting CellDyc-inferred transcriptomic velocities onto the original UMAP embedding, we faithfully recapitulated the erythroid differentiation trajectory. Unexpectedly, this analysis identified a reversed trajectory segment within Blood Progenitor 2 (BP2) cells (Fig. 5b), suggesting that erythropoiesis involves state transitions more complex than a simple unidirectional differentiation process. Analysis of G2M scores, calculated using the scVelo package [27], further revealed a progressive decline in self-renewal capacity throughout erythropoiesis (Fig. 5c), consistent with previous reports [50].

**Figure 5.**
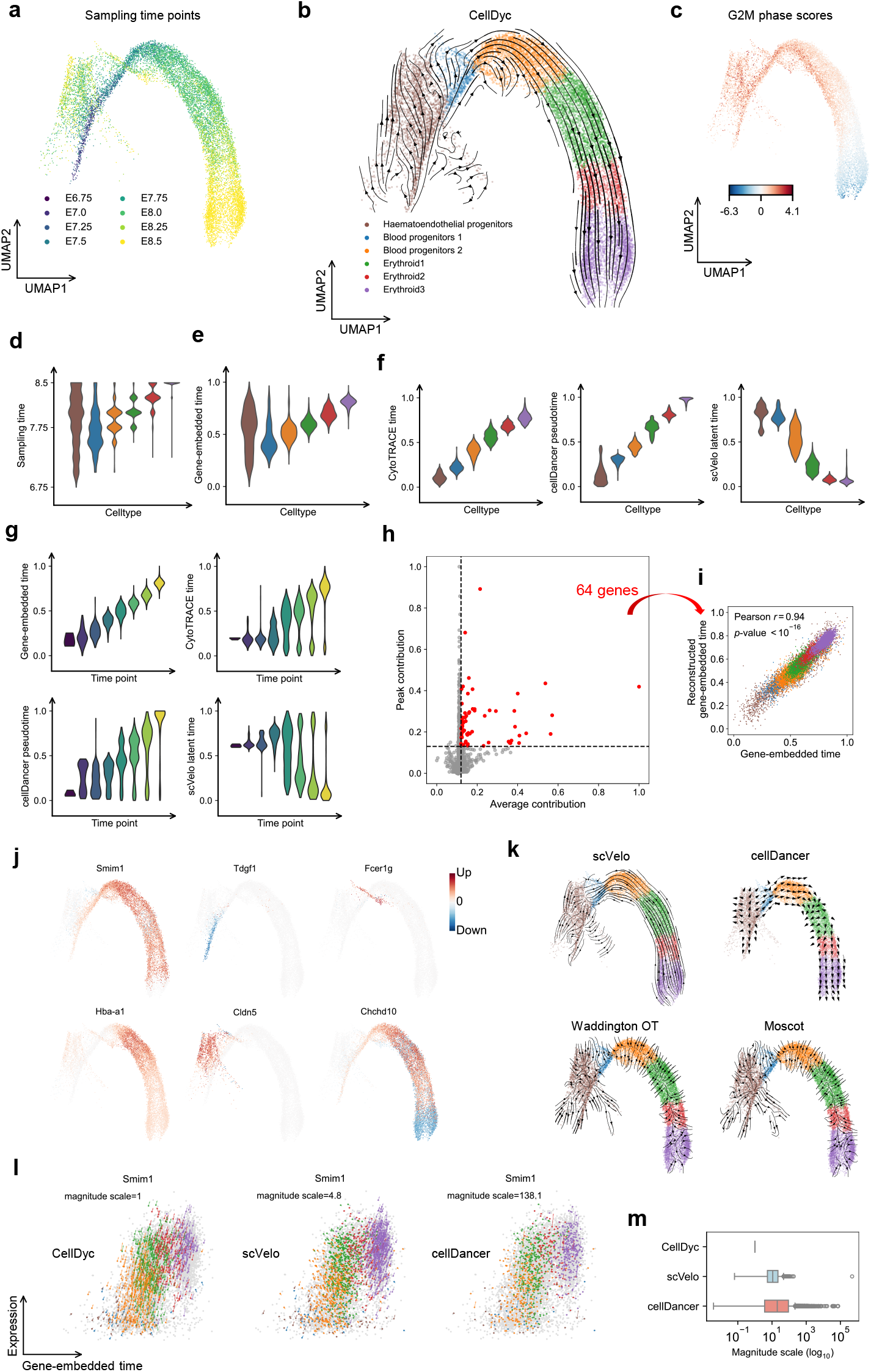
Reconstruction of the gastrulation erythroid maturation trajectory and recovery of inherent temporal heterogeneity. **a**, UMAP embedding of 12324 cells from the process of gastrulation erythroid maturation. Cells are colored according to sampling time points. **b**, Transcriptomic velocities inferred by CellDyc were projected onto UMAP in (**a**), faithfully recapitulating the erythroid differentiation trajectory and unexpectedly revealing a reversed trajectory segment within Blood Progenitor 2 (BP2) cells. **c**, UMAP embedding as in (a), colored by G2M phase scores to indicate self-renewal capacity. **d**, Violin plots of observed sampling time across cell types. **e**, Violin plots of predicted gene-embedded time across cell types. **f**, Violin plots of time predicted by different methods across cell types. Left: CytoTRACE time; Middle: cellDancer pseudotime; Right: scVelo latent time. **g**, Violin plots of time predicted by different methods across sampling time points.(Upper left) CellDyc gene-embedded time; (Upper right) CytoTRACE time; (Lower left) cellDancer pseudotime; (Lower right) scVelo latent time. **h**, Scatter plot of average and peak contributions to gene-embedded time for individual genes. Each dot represents a single gene (x-axis: average contribution; y-axis: peak contribution). Genes with high contributions (average > 0.12 and peak > 0.13) are highlighted (n = 64). **i**, Scatter plot of reconstructed versus original gene-embedded time based on 64 genes with high contributions (as in (**h**)), colored by cell types. Pearson’s correlation coefficient is indicated. **j**, Heatmap of single-gene velocities. Red indicates instantaneous increase in expression; blue indicates instantaneous decrease. **k**, Projection of RNA velocities or cell state transitions derived from different methods onto UMAP. Top left: scVelo. Top right: cellDancer. Bottom left: Waddington-OT. Bottom right: Moscot. **l**, Velocity dynamics of Smim1 inferred by different methods. Expression levels (y-axis) are plotted against gene-embedded time (x-axis). Arrows depict the velocities scaled by the indicated magnitude scale. The magnitude scale represents the scaling factor required to align the velocities with the gene-embedded time for each respective method. Methods presented (from left to right): CellDyc, scVelo, and cellDancer. **m**, Boxplots of magnitude scales for aligning per-gene velocities from different methods with gene-embedded time.

This developmental complexity manifests as pronounced temporal heterogeneity across lineages. Examination of the distributions of experimental time revealed that the temporal span represented by each cell type progressively contracted from hematoendothelial progenitors to BP2 cells (Fig. 5d), mirroring the observed decline in self-renewal capacity (Fig. 5c). Notably, in hematoendothelial progenitors, self-renewal potential was not reflected through a dominant cell cycle trajectory, but rather through gradual yet directional state changes spanning from E6.75 to E8.5 (Fig. 5a). These findings underscore that distinct cell populations exhibit markedly different rates of state change during differentiation.

The CellDyc-derived gene-embedded time accurately recapitulated experimental time, preserving both the sequential order of cell type emergence and the specific temporal windows occupied by each population (Fig. 5e). In contrast, alternative methods exhibited substantial limitations: CytoTRACE time (1 - CytoTRACE score [48]) and cellDancer velocity pseudotime [28] captured inter-cell-type temporal relationships but failed to preserve the temporal span of individual cell types. Most critically, scVelo latent time [27] failed to reconstruct even basic temporal relationships due to erroneous inference of differentiation directionality (Fig. 5f, 5k). Tests regarding the recapitulation of experimental time points yielded consistent results (Fig. 5g).

To elucidate the mechanistic basis of this temporal encoding, we identified 64 pivotal time-encoding genes based on their average and peak contributions to gene-embedded time (Fig. 5h, Methods). Remarkably, direct reconstruction using only these genes, weighted by their Gene Clock contributions, nearly perfectly recapitulated the original gene-embedded time (Fig. 5i). The Gene Clock integrates genes exhibiting diverse expression dynamics (Fig. 5j): *Smim1* showed near-universal upregulation; *Tdgf1, Fcer1g*, and *Cldn5* each displayed monotonic changes during specific developmental windows, enabling the capture of cell-type-specific temporal dynamics; *Hba-a1* captured the rapid transition from BP2 to Erythroid 3; and *Chchd10* demonstrated that temporal encoding extends beyond monotonically regulated genes.

Our comparative analyses revealed substantial discrepancies between RNA velocity-derived temporal estimates and experimental time (Fig. 5d, 5f). Since velocity theoretically represents the time derivative of gene expression, these discrepancies indicate the presence of previously unrecognized errors stemming from deficiencies in temporal model fitting. We therefore computed a magnitude scale for each gene, quantifying the scaling factor required to align expression time derivatives with gene-embedded time (Methods). We visualized the velocities of *Smim1* from CellDyc, scVelo [27], and cellDancer [28] in the Expression-Time space after scaling. CellDancer’s predictions showed generally correct directional alignment, whereas scVelo’s were predominantly incorrect—a pattern consistent with their macroscopic trajectory prediction performance (Fig. 5k, 5l). Notably, aligning *Smim1* predictions required a 4.8× magnification for scVelo, whereas cellDancer required a substantial 138.1× magnification to achieve proper Expression-Time space representation (Fig. 5l).

Examination of magnitude scale distributions across all predictive genes revealed that scVelo’s scales spanned approximately one order of magnitude, whereas cellDancer’s spanned approximately two orders of magnitude (Fig. 5m). This difference reflects their distinct design philosophies: scVelo explicitly models latent time and performs per-gene scale correction based on gene-specific timing [27], whereas cellDancer lacks explicit temporal encoding and cannot implement such corrections [28]. Additional Expression-Time plots demonstrated that both scVelo and cellDancer exhibited not only substantial inter-gene magnitude scale variability but also intra-gene heterogeneity in arrow lengths compared to CellDyc (Extended Data Fig. 5). These variations reflect a fundamental temporal misalignment that persists even after cross-gene scale correction, and establish CellDyc as a robust reference for benchmarking and improving velocity-based methods.

### CellDyc Enables Accurate Fate Probability Inference in Cellular Reprogramming

We applied CellDyc to a reprogramming dataset comprising six experimental time points (Fig. 6a). In this study, Biddy et al. performed direct lineage reprogramming of mouse embryonic fibroblasts (MEFs) to induced endoderm progenitors (iEPs) [11], employing the CellTagging lineage tracing technique to label cells traversing distinct trajectories—one leading to successful reprogramming and another to a 'dead-end’ state—and annotated clusters corresponding to different terminal fates (Fig. 6b). To evaluate whether CellDyc could accurately delineate individual cell fates, we selected 3,049 lineage-barcoded cells as a benchmark. While CellDyc focuses on predicting transcriptomic velocities, its outputs are designed for seamless integration with existing computational infrastructure; here, we employed CellRank [43, 44] to compute fate probabilities based on the transcriptomic velocities inferred by CellDyc.

**Figure 6.**
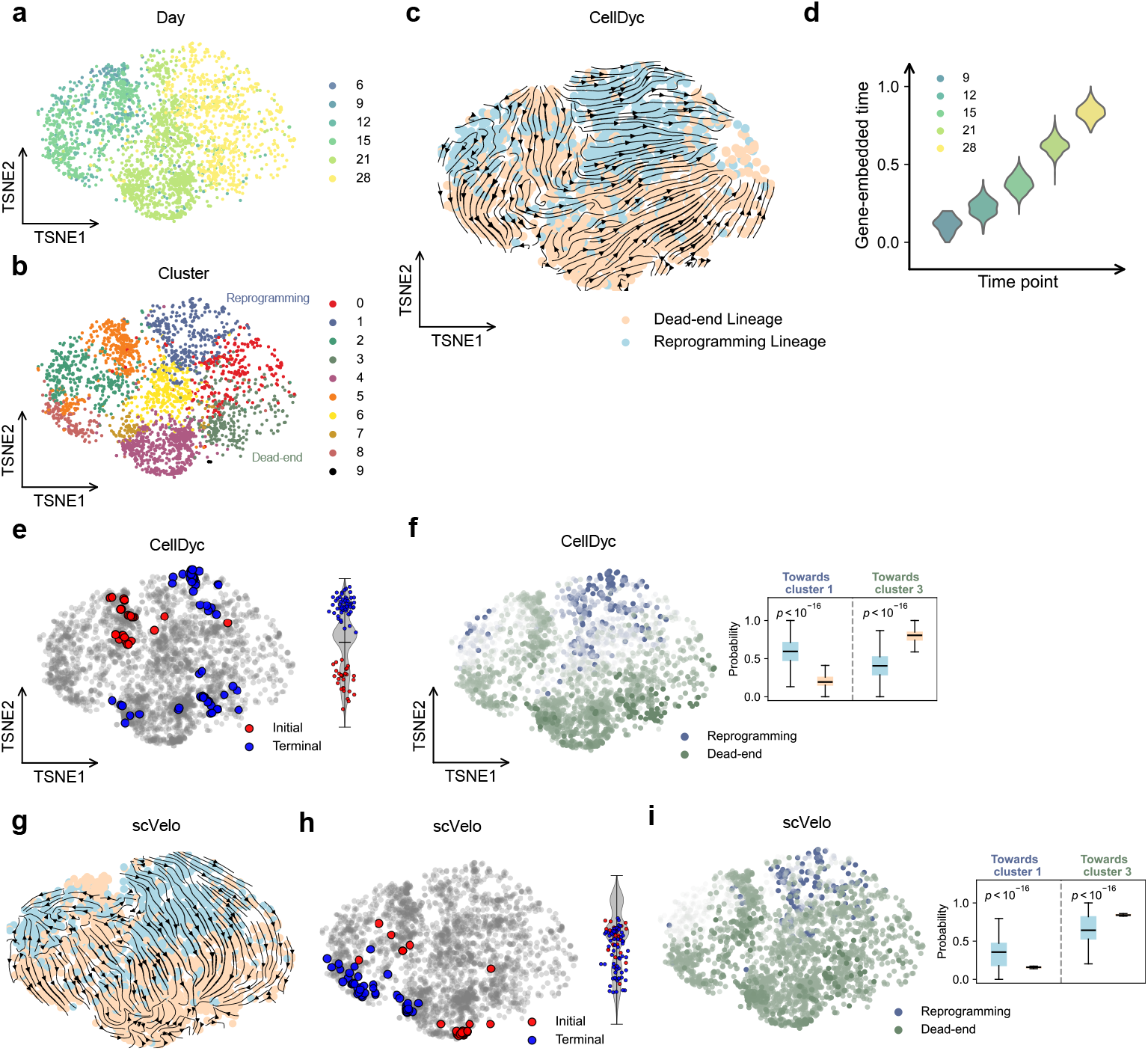
CellDyc-guided analysis of cell fate probabilities in mouse embryonic fibroblast reprogramming. **a**, t-SNE embedding of 3,049 mouse embryonic fibroblasts (MEFs) reprogramming to induced endoderm progenitors (iEPs). Cells are colored by day after reprogramming induction. **b**, The same t-SNE embedding as in (**a**), but colored by cluster annotations. Clusters 1 and 3 represent the successful reprogramming trajectory and the dead end, respectively. **c**, Transcriptomic velocities derived from CellDyc were projected onto t-SNE in (**a**). Cells are colored by lineage barcodes from CellTagging lineage tracing. **d**, Violin plots of predicted gene-embedded time across sampling time points measured in days after reprogramming induction (Day 6, containing only three cells, is not shown). **e**, The initial and terminal macrostates were determined using CellRank based on transcriptomic velocities derived from CellDyc. In the left panel, the initial states are marked as red dots and the terminal states are marked as blue dots on the t-SNE. The right panel shows the distribution of cells belonging to the initial and terminal macrostates across gene-embedded time, demonstrating a clear temporal separation between initial and terminal cells. **f**, Cell fate probability to cluster 1 versus cluster 3. Left: t-SNE projection showing the computed probability of each cell reaching the combined macrostates of cluster 1 and cluster 3; colors follow the same fate palette as in panel (**b**), with darker shades indicating higher probability. Right: Boxplots of fate probabilities for cells from Reprogramming Lineage and Dead-end Lineage, showing Reprogramming Lineage cells are significantly enriched for the cluster 1 fate while Dead-end Lineage cells are enriched for the cluster 3 fate (one-sided Mann-Whitney U test, *p* < 1.0 × 10^−16^ for both clusters). **g**, Velocities derived from scVelo projected onto the t-SNE in (**a**). Cells are colored by lineage barcodes from CellTagging lineage tracing. **h**, Similar analysis to (**e**) but with velocities derived from scVelo. The left panel visualizes the initial (red) and terminal (blue) macrostates on the t-SNE. The right panel displays these cells across gene-embedded time, revealing that the initial and terminal states are intermingled rather than temporally separated. **i**, Cell fate probabilities computed using velocities from scVelo with manually anchored terminal states. The left panel displays the t-SNE projection of probabilities for cluster 1 and cluster 3. The right panel shows boxplots of fate probabilities stratified by lineage, indicating that, when terminal states are constrained to match those in the CellDyc analysis, the distinct enrichment of Reprogramming Lineage cells toward cluster 1 and Dead-end Lineage cells toward cluster 3 is preserved (one-sided Mann-Whitney U test, *p* < 1.0 × 10^−16^ for both clusters).

Projection of CellDyc-derived transcriptomic velocities onto the original t-distributed stochastic neighbor embedding (t-SNE) revealed discernible directional flows from day 6/9 cells toward the two terminal states (clusters 1 and 3) (Fig. 6c). Furthermore, the inferred gene-embedded time effectively reconstructed the experimental time (Fig. 6d; day 6 contained only three cells and is not shown). Given that cell lineages corresponding to the two fates were intermingled in the low-dimensional embedding, our primary focus was the accuracy of fate assignment. We first identified initial and terminal macrostates using CellRank based on CellDyc’s predictions. A subset of cluster 5 was identified as the initial state, while clusters 1 and 3 represented the primary terminal states. Notably, the distribution of gene-embedded time for initial versus terminal cells exhibited a clear temporal separation (Fig. 6e). Using all macrostates associated with clusters 1 and 3 as the two designated terminal states, we computed fate probabilities for each cell. Cells from the Reprogramming Lineage showed significant enrichment for the cluster 1 fate (one-sided Mann-Whitney U test, p < 1.0 × 10^−16^), whereas Dead-end Lineage cells were strongly enriched for the cluster 3 fate (one-sided Mann-Whitney U test, p < 1.0 × 10^−16^) (Fig. 6f).

In contrast, the projection of transcriptomic velocities derived from scVelo [27] failed to reveal clear paths toward either terminal state (Fig. 6g). When used as input for CellRank, scVelo incorrectly identified initial states primarily within cluster 1 and terminal states mainly in cluster 4. Critically, the gene-embedded time distributions for these scVelo-identified initial and terminal cells were overlapping and intermingled rather than temporally distinct (Fig. 6h). However, when we manually constrained the terminal states to match those identified in the CellDyc analysis, scVelo was then able to correctly enrich cells from each lineage in their respective terminal fates (Fig. 6i). Similarly, OT-based methods Waddington-OT [39] and Moscot [40] identified incorrect initial states and failed to automatically detect coherent terminal states (Extended Data Fig. 6c-d). While Waddington-OT could produce correct fate probability predictions when provided with the same manually anchored terminal states (Extended Data Fig. 6e), Moscot failed to simultaneously align clusters 1 and 3 during macrostate construction, precluding valid fate probability analysis. These results demonstrate that the automatic and accurate identification of initial and terminal states depends critically on the correct directionality of transcriptomic velocities. Yet, once appropriate terminal states are specified, fate probability assignments may not be solely determined by velocity vectors but are also influenced by transcriptomic similarity.

### CellDyc Generalizes to Inferred Time and Rectifies Temporal Distortion to Recover Transcriptomic Velocity

To assess the generalization capability of CellDyc beyond experimental time points, we evaluated whether the framework could utilize inferred temporal coordinates—specifically pseudotime and CytoTRACE time (1-CytoTRACE score [48]). While existing tools like CellRank [44] have introduced specific kernels (PseudotimeKernel and CytoTRACEKernel) to directly infer cell state transitions from such metrics, these inputs represent relative orderings rather than physical time, often posing challenges for resolving cellular dynamics. Proceeding from our premise that gene-embedded time is an intrinsic property of the transcriptome, we posited that CellDyc could leverage these inferred temporal proxies as weak supervision to uncover the objective, latent temporal progression embedded within gene expression patterns and effectively infer transcriptomic velocity.

We utilized the zebrafish embryogenesis dataset (Fig. 3h) to test the generalization capability. First, we calculated pseudotime using the diffusion pseudotime (DPT) algorithm as implemented in Scanpy [20, 51], initialized from the 3.3-hour developmental stage (Fig. 7a). Although the pseudotime generally recapitulated the sequential order of development, the predicted intervals displayed significant distortion relative to the actual experimental sampling times (Fig. 7b). Despite this non-linear distortion in the input labels, CellDyc successfully reconstructed transcriptomic velocity vector fields that aligned well with the known developmental trajectory (Fig. 7c). Strikingly, the gene-embedded time recovered by CellDyc exhibited a higher correlation with the experimental time than the original pseudotime input used for training (Fig. 7d). This suggests that the model effectively “rectified” the temporal distortion inherent in the pseudotime labels. In contrast, CellRank failed to establish a coherent trajectory using the PseudotimeKernel on this dataset (Fig. 7e).

**Figure 7.**
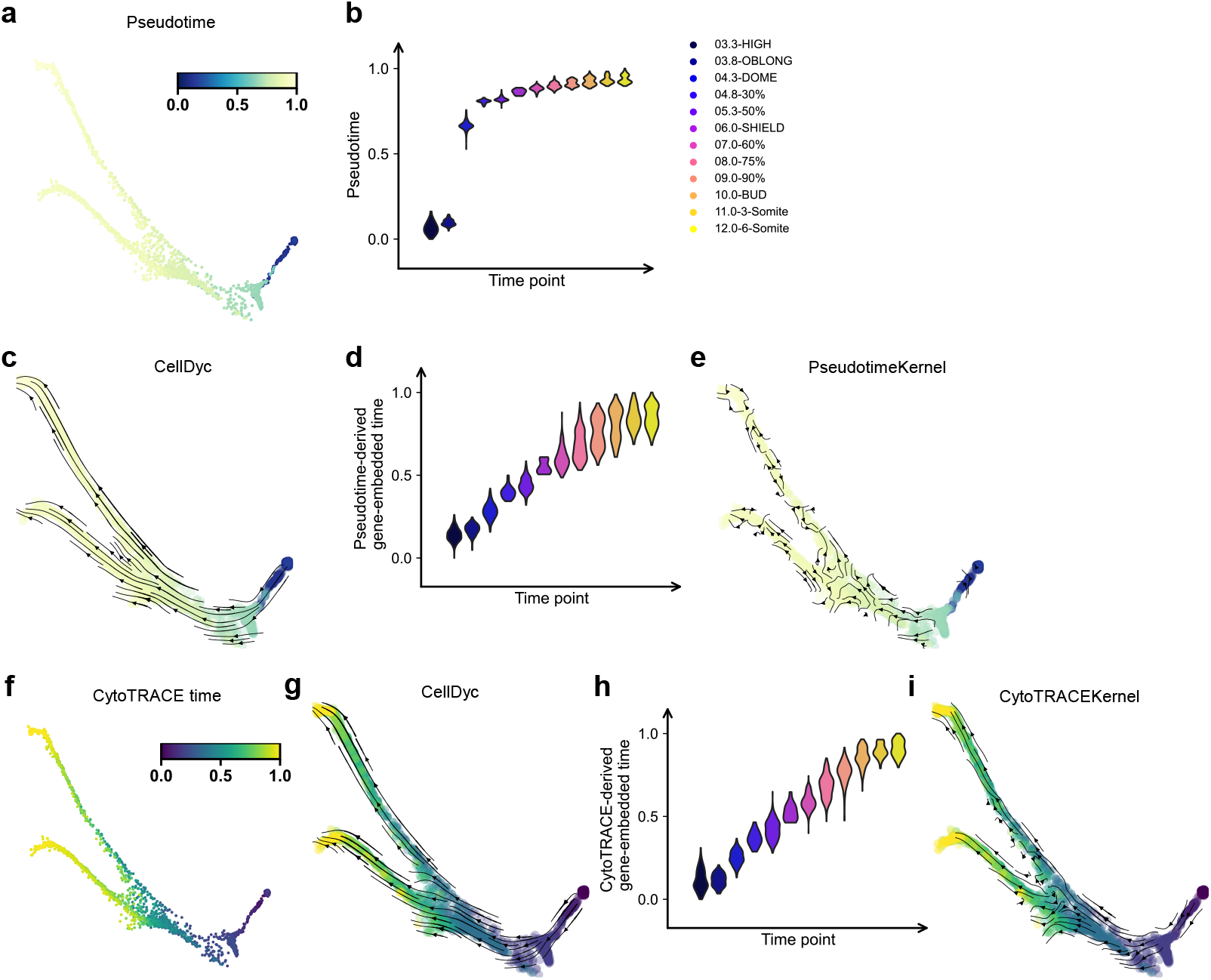
Recovery of transcriptomic velocity and gene-embedded time using inferred temporal priors. **a**, Force-directed layout of zebrafish embryogenesis (as in Fig 3h). Cells are colored by pseudotime. **b**, Violin plots of pseudotime across sampling time points. **c**, Projection of transcriptomic velocities derived from CellDyc (trained with pseudotime) onto the force-directed layout. **d**, Violin plots of predicted gene-embedded time across sampling time points from CellDyc trained with pseudotime. **e**, Projection of cell state transitions derived from the PseudotimeKernel of CellRank (using pseudotime) onto the force-directed layout. **f**, Force-directed layout of zebrafish embryogenesis (as in Fig. 3h). Cells are colored by CytoTRACE time. **g**, Projection of transcriptomic velocities derived from CellDyc (trained with CytoTRACE time) onto the force-directed layout. **h**, Violin plots of predicted gene-embedded time across sampling time points from CellDyc trained with CytoTRACE time. **i**, Projection of cell state transitions derived from the CytoTRACEKernel of CellRank (using CytoTRACE time) onto the force-directed layout.

We subsequently evaluated CytoTRACE time (1 - CytoTRACE score), which inherently exhibited a temporal distribution closer to the experimental time points than pseudotime did (Fig. 7f,Fig. 3j). Consistent with previous results, CellDyc trained on CytoTRACE time inferred transcriptomic velocities that accurately captured the developmental flow (Fig. 7g) and yielded a gene-embedded time metric that was again more consistent with experimental time than the input labels (Fig. 7h). While CellRank’s performance improved significantly with the CytoTRACEKernel compared to its performance with pseudotime (Fig. 7i), it still did not achieve the trajectory resolution provided by CellDyc.

The remarkable ability of CellDyc to recover time distributions consistent with experimental ground truth—even when trained on highly distorted pseudotime inputs—underscores a fundamental insight: gene-embedded time is an objective reality inherent to the transcriptomic data. In this framework, external temporal labels serve primarily as a guide to align the model with this intrinsic coordinate system. Consequently, CellDyc does not merely fit the input labels; rather, it mines latent temporal information to rectify input bias, thereby recovering a temporal metric closer to the “real-world” biological process. Theoretically, CellDyc is capable of accepting any continuous temporal annotation, demonstrating its potential as a flexible tool that can be integrated into diverse computational workflows to translate any temporal proxies into high-resolution transcriptomic velocities.

## Discussion

Reconstructing continuous cellular dynamics from time-series scRNA-seq snapshots remains a fundamental challenge in computational biology. Historically, the field has faced a dichotomy: Optimal Transport (OT) methods [33-40] leverage experimental time points to ensure global directionality but typically treat these time points as static distributions, thereby lacking local resolution and ignoring dynamics occurring within time points. Conversely, RNA velocity [26-30] provides instantaneous transcriptomic velocities but generally disregards explicit temporal information, limiting it to splicing kinetic assumptions and rendering it susceptible to stochastic noise. CellDyc resolves this trade-off by introducing a semi-supervised learning paradigm that effectively bridges the gap between macroscopic trends and microscopic cellular dynamics. Unlike OT-based approaches that assume cell state transitions occur solely between discrete time points, CellDyc’s strategy applies supervision only where time labels clearly indicate transient dynamics. This allows the model to learn generalizable laws of state evolution, enabling the prediction of vector fields even in the “unlabeled intervals” within time points and robustly recovering continuous dynamics both across and within snapshots.

Our findings suggest that gene-embedded time is an intrinsic biological feature to be uncovered. It is neither a synthetic metric to be invented, nor a simple interpolation of experimental time points, but rather a biologically grounded temporal coordinate system naturally encoded within the transcriptome. Our results demonstrate that this intrinsic timeline can be recovered using various forms of supervision labels—ranging from discrete experimental time points to inferred pseudotime. Crucially, the fidelity of this recovery relies not merely on a specialized loss function design, but on the synergistic interplay between CellDyc’s two prediction tasks. By analyzing the model’s dual-loss architecture, we found that the time prediction loss serves as a scaffold, designating a latent space for explicit temporal encoding, while the trend prediction task refines this space concurrently with the inference of transcriptomic velocities. This synergy allows CellDyc to overcome the limitations of its supervision signals—effectively “denoising” technical artifacts (e.g., in Zman-seq [42]) or correcting distortions in pseudotime—to reconstruct a metric that more closely approximates the true physical progression of the system.

The biological utility of this intrinsic temporal coordinate is profound. By establishing this stable intrinsic coordinate system, CellDyc is empowered to delineate subtle variations in cellular behavior that are otherwise obscured. In the glioblastoma microenvironment, we identified that the immunosuppressive microenvironment delays the differentiation of monocytes, a quantitative insight inaccessible to standard trajectory inference. Similarly, in mouse gastrulation, CellDyc resolved a progressive contraction of the temporal span from progenitors to differentiated states, revealing lineage-specific temporal heterogeneity during erythroid maturation. These findings underscore that gene-embedded time provides a rigorous quantitative basis for comparing the “tempo” of cellular life across different conditions and lineages.

Furthermore, CellDyc is designed to serve as a versatile hub within the current computational ecosystem. It functions as a bridge, accepting upstream inputs ranging from raw experimental time to inferred times (e.g., pseudotime [20-25], CytoTRACE [48]) and converting them into vector-based transcriptomic velocities. For downstream analysis, CellDyc seamlessly interfaces with frameworks originally designed for RNA velocity, such as PAGA [52], CellRank [43, 44], and dynamo [53]. As demonstrated in our fibroblast reprogramming study, CellDyc-derived velocities enabled CellRank to correctly map initial and terminal fates without relying on splicing models, succeeding where traditional RNA velocity failed due to technical noise.

However, our approach is not without limitations, which point toward important avenues for future development. The current CellDyc framework operates under a “transcriptome-centric” assumption—that the transcriptomic state alone is sufficient to describe cell fate. This view may oversimplify scenarios driven by epigenetic priming [54], chromatin remodeling [55], or translational regulation [56]. This limitation, however, highlights the potential for extending CellDyc to multi-omics data. Unlike RNA velocity methods bound by specific ordinary differential equation (ODE) models [26-29, 57] of splicing kinetics, CellDyc’s data-driven framework can be readily adapted to learn “multi-modal velocities” from platforms like CITE-seq or 10x Multiome [58-60]. Additionally, the rise of spatiotemporal transcriptomics offers a new frontier. While recent OT-based methods have begun to explore dynamics across spatial slices [61], they share the limitation of ignoring intra-slice dynamics. Extending CellDyc to incorporate spatial coordinates could integrate cell-cell interactions [62-64] into velocity predictions, paving the way for a comprehensive deciphering of how microenvironmental cues drive cell fate decisions in both time and space.

## Acknowledgements

This work was supported by grants from National Natural Science Foundation of China (32341007), and the Fundamental Research Funds for the Central Universities (020814380190, 020814380152).

## Author contributions

X.Z. and X.S. conceived the study. X.Z. designed the method and implemented the software. X.S. performed model validation and conducted most analyses. N.Z., Y.W., M.D., and X.L.Z. contributed to data collection, discussion, and software testing. J.Y. and C.-Y.Z provided constructive suggestions. X.W., Z.Z., and J.W. supervised the study and provided constructive suggestions. X.Z. and X.S. wrote the manuscript with contributions from all co-authors.

## Competing interests

The authors declare that a patent application related to this work has been filed, listing X.Z., X.S., J.Y., Z.Z., and J.W. as inventors.

## Data availability

All the scRNA-seq raw data are publicly accessible. The *C. elegans* embryonic data are available under GEO accession number GSE126954; the analysis-ready data in.h5ad format are also provided at https://zenodo.org/records/7496490. The zebrafish embryogenesis data subset used in this study can be extracted using CellRank’s CLI: cellrank.datasets.zebrafish(), with the full raw dataset archived under GEO accession number GSE106587. The raw data for the glioblastoma-associated monocyte differentiation are available under GEO accession number GSE232040. The mouse gastrulation erythropoiesis data can be extracted using scVelo’s CLI: scvelo.datasets.gastrulation() or from ArrayExpress under accession number E-MTAB-6967. The mouse embryonic fibroblast reprogramming data with lineage information can be extracted using CellRank’s CLI: cellrank.datasets.reprogramming_morris (subset='48k’), with the raw data available under GEO accession number GSE99915. All processed data used in this study are available for download from our Zenodo repository at https://zenodo.org/records/18639013.

## Code availability

CellDyc is released under the BSD-3-Clause license, with the source code available at https://github.com/hsinring/celldyc. Code to reproduce the results in this paper can be found at https://github.com/hsinring/celldyc/reproducibility.

## Methods

### CellDyc Architecture

We propose a semi-supervised deep learning framework to infer transcriptomic velocity and gene-embedded time from single-cell transcriptome profiles. The model accepts a gene expression vector **x** ∈ ℝ^*g*^, where g denotes the number of genes. The architecture comprises a linear embedding layer followed by a multi-head decoder with a shared backbone network (Fig. 1b). The forward pass is defined by the following sequential transformations:

#### Linear embedding

The input transcriptome profile is projected into a low-dimensional latent space:

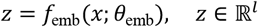

where *l* is the latent dimensionality (default *l* = 10).

#### Gene-embedded time extraction

Temporal information is explicitly encoded in the first latent dimension. Gene-embedded time *ĉ* is defined as:

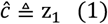

#### Shared backbone network

The latent representation ***z*** is transformed by a fully-connected backbone network into a hidden representation:

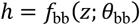

#### Non-steady-state probability head

This output head predicts, for each gene, the probability of being in a dynamic, non-steady-state regime:

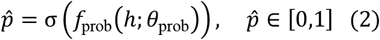

where σ(·) denotes the logistic sigmoid function.

#### Expression trend head

This head predicts the probability of directional expression changes, with outputs centered at zero to enable symmetric representation of up- and down-change:

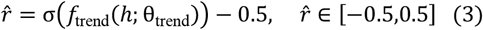

#### Model parameterization

The complete model is parameterized by the set *θ* = {*θ*_emb_, *θ*_bb_, *θ*_prob_, *θ*_trend_}, corresponding to the embedding layer, backbone network, probability head, and trend head, respectively.

### Semi-supervised Training

#### Temporal Pairwise Ranking Loss

Experimental time points typically provide only coarse temporal relationships between cellular samples, without revealing fine-grained temporal ordering within each time point. Direct regression on discrete time points is insufficient to capture the underlying continuous processes. As illustrated in Fig. 1b, converting discrete time points into pairwise rank relationships between individual cells yields comprehensive supervisory signals for the majority of cell pairs, enabling a semi-supervised time representation learning framework. Consider a mini-batch of *b* cells, where each cell *i* is associated with an observed time point *t*_*i*_ and a predicted time representation *Ĉ*_*i*_ learned by the model (as shown in equations (1)). We construct pairwise difference matrices to capture relative temporal relationships:

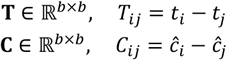

where **T** encodes the ground-truth temporal differences, and **C** represents the corresponding differences in the predicted time representation.

To eliminate invalid comparisons between cells derived from the same experimental time point, we introduce a binary masking matrix:

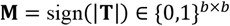

where *M*_*ij*_ = 1 if |*t*_*i*_ − *t*_*j*_ | > 0, and 0 otherwise. This mask ensures that the loss is computed exclusively over inter-time-point pairs, where a meaningful temporal ordering exists.

The temporal loss ℒ_time_ enforces directional consistency between the predicted-based differences and the ground-truth temporal differences. For each cell *i*, we maximize the cosine similarity between the masked predicted difference vector and the temporal difference vector:

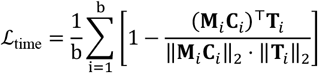

where **M**_*i*_, **C**_*i*_and **T**_*i*_ denote the *i*-th rows of **M, C** and **T** respectively, and ‖ ⋅ ‖_2_ denotes the *ℓ*_2_ norm. The inner term computes the masked cosine similarity for cell *i*, and the loss penalizes deviations from perfect alignment. This formulation is fully differentiable and operates on pairwise relationships without requiring explicit negative sampling.

This loss function possesses several desirable characteristics: (1) it is invariant to the absolute scale of the time representation, focusing instead on preserving relative temporal ordering; (2) it automatically down-weights large-magnitude differences, preventing a small subset of pairs with extensive temporal gaps from disproportionately dominating the loss function and thereby enhancing model robustness; and (3) it enables semi-supervised learning by leveraging the structure inherent in the temporal data itself. The resulting supervision signal drives the model to learn a continuous, biologically meaningful time representation.

#### Trend Alignment Loss

To enhance effective supervision for trends prediction task, we implemented a neighborhood-based temporal smoothing procedure. For each cell, we computed its local neighborhood using Scanpy’s neighbor detection algorithm [51]. Following [65, 66], within each neighborhood, we smoothed the time point annotation by transforming the temporal distribution into a cumulative distribution area-under-the-curve (AUC) metric, but we designed a novel and more efficient algorithm for its implementation.

Assuming the time point annotation contains *k* + 1 ordered time points *t*_0_ < *t*_1_ < *t*_2_ < ⋯ < *t*_*k*_, we first converted each discrete time point t_m_ (where *m* = 0, 1, …, *k*) into its normalized area contribution within the cumulative distribution:

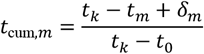

where the interval adjustment term δ_m_ accounts for half the distance between consecutive time points:

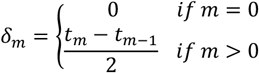

This transformation maps each time point to a normalized contribution value *t*_cum,*m*_ ∈ [0,1].

For a given cell *i* with *n*_neighbors_ cells in its neighborhood 𝒩_*i*_, we computed the smoothed temporal value *t*_smooth, *i*_ as:

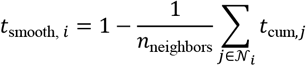

where *t*_cum,*j*_ denotes the area contribution of neighbor cell *j*. This formulation derives the smoothed time from the mean area contribution of neighboring cells, which represents the AUC of cumulative temporal distribution.

For scalable application to all *n* cells in the dataset, we implemented a matrix-based computational scheme. Let **A** ∈ ℝ^*n*×*n*^ be the connectivity matrix derived from Scanpy’s neighborhood graph, where element *A*_*ij*_ ≥ 0 represents the connection weight between cell *i* and cell *j* (with *A*_*ij*_ = 0 indicating no neighborhood relationship). Let **T**_cum_ ∈ ℝ^*n*^ be the vector of all cells’ area contributions. The smoothed temporal values for all cells **T**_smooth_ ∈ ℝ^*n*^ were then calculated as:

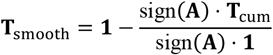

where sign(**A**) binarizes the connectivity matrix (1 for connected neighbors, 0 otherwise), ⋅ denotes matrix multiplication, and **1** is a column vector of ones. This formulation efficiently computes, for each cell, one minus the mean *t*_cum_ value across its neighborhood, enabling rapid processing of large-scale datasets.

To supervise the training of trends prediction, we developed a loss function that leverages temporal ordering and local transcriptomic similarity. For each cell, a ground-truth directional vector and a reliability score vector are first computed from the data; these are then compared against model predictions via a weighted cosine distance.

Let 𝒩_𝒾_ denote the neighborhood of cell i, which contains *n*_neighbors_ cells. Denote by **x**_*i*_, **x**_*j*_ ∈ ℝ^*g*^ the g-dimensional gene-expression profiles of cell *i* and neighbor*j* ∈ 𝒩_𝒾_, respectively, and by *t*_*smooth,i*_, *t*_*smooth,j*_ their corresponding smoothed time values. The ground-truth directional vector *r*_*i*_ ∈ ℝ^*g*^ encodes the expected per-gene expression change for cell *i* and is calculated as:

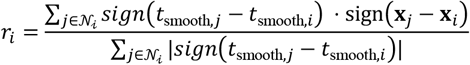

Here, the temporal weight sign (t_smooth,*j*_ − t_smooth,i_) assumes a value of +1 if neighbor *j* lies in the future (i.e., has a larger time), −1 if it lies in the past, and 0 if the times are equal. The term sign (**x**_*j*_ − **x**_*i*_) denotes the element-wise sign of the expression difference vector, encoding the direction of change for each gene. The numerator aggregates the transcriptomic shift directions of all neighbors, weighted by their temporal relationship to cell *i*. The denominator serves as a normalization factor equal to the number of neighbors with non-identical time values, thereby excluding temporally coincident cells. Each entry of *r*_*i*_ thus lies in the interval [−1,1] and represents the predicted direction of expression change for the corresponding gene as a function of time.

The reliability score vector ρ_i_ ∈ ℝ^*g*^ measures the temporal dispersion within 𝒩_𝒾_ and is defined as:

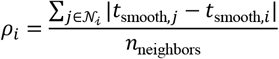

A larger *ρ*_*i*_ indicates that the neighborhood spans a broader range of time, implying a more robust temporal signal for inferring transcriptional dynamics.

Let 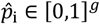 denote the model-predicted probability that each gene of cell *i* resides in a dynamic, non-steady-state regime, and let 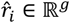 denote the model-predicted directionality vector (as shown in equations (2)(3)). The ground-truth trend vector is *r*_*i*_, with associated reliability *ρ*_*i*_ (computed above). The trend alignment loss is defined as a reliability-weighted cosine distance:

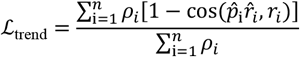

where *n* is the total number of cells and cos(⋅,⋅) denotes cosine similarity:

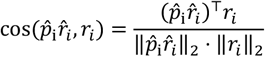

This formulation ensures that each cell contributes proportionally to its reliability, with the denominator normalizing by the total reliability across all cells to yield a weighted average of directional prediction errors.

#### Training Procedure

The learning process of CellDyc is mathematically formulated as an optimization problem aimed at capturing both gene-embedded time and the directionality of expression changes. The training objective is defined by the following composite loss function:

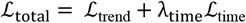

where λ_time_ (default: 0.1) represents a manually tuned hyperparameter that modulates the balance between trend preservation and temporal consistency. The systematic impact of this hyperparameter on model convergence and performance is investigated in Fig. 4m.

During training, we minimize ℒ_total_ using the AdamW [67]optimizer with a learning rate of × 10^−2^ and weight decay of 1 × 10^−2^. The learnable parameter ensemble *θ* = {*θ*_emb_, *θ*_bb_, *θ*_prob_, *θ*_trend_} is iteratively updated through backpropagation.

### Transcriptomic Velocity Prediction

Following semi-supervised training, CellDyc provides for each cell *i* with gene expression vector **x**_*i*_ ∈ ℝ^*g*^ three key outputs: a time representation *Ĉ*_*i*_, a non-steady probability vector 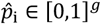, and a directionality vector 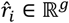 (Equations (1)-(3)). However, the complete transcriptomic velocities (*d***x**/*dĈ*)_*i*_ ∈ ℝ^*g*^ remains partially uncharacterized. While 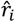 reliably indicates the directional change for each gene, the element-wise magnitude of transcriptomic velocities ‖(*d***x**/*dĈ*)_*i*_ ‖ is not yet quantified.

#### Magnitude Quantification

we compute the magnitude vector through a neighborhood-based standardization approach. Let 𝒩_𝒾_ denote the neighborhood of cell *i*, comprising *n*_neighbors_ cells, where each neighbor *j* ∈ 𝒩_𝒾_ is characterized by its gene expression vector **x**_*j*_ ∈ ℝ^*g*^ and inferred time *Ĉ*. Let *s*_*i*_ (⋅) represent the element-wise sample standard deviation operator computed over 𝒩_𝒾_.

The magnitude of transcriptomic velocities for cell *i* is defined as:

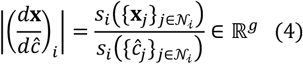

where the numerator is computed element-wise across genes:

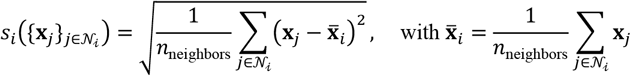

and the denominator is a scalar representing the standard deviation of temporal coordinates within the neighborhood:

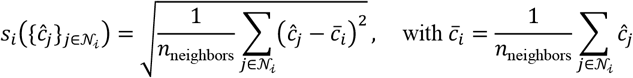

#### Processing of Probability and Directionality Vectors

Prior to velocities calculation, the predicted vectors undergo threshold-based filtering using a hyperparameter threshold *τ* (default: 0.05). The non-steady probability vector 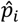 is binarized to retain only high-confidence predictions:

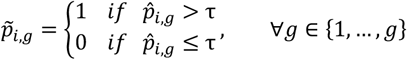

Concurrently, the directionality vector 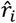 is converted to its sign representation:

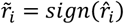

The directional indicator ***m***_*i*_ is then derived via element-wise multiplication:

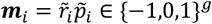

This ensures only genes with sufficient non-steady probability contribute directional information to the final velocities prediction.

#### Final Transcriptomic Velocity Prediction

Integrating the filtered directional indicator with the magnitude yields the complete transcriptomic velocities:

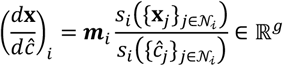

This final formulation provides a comprehensive quantification of transcriptomic velocities that incorporates time representation *Ĉ*_*i*_, distinguishes steady from non-steady genes by filtering low-confidence predictions via the hyperparameter *τ*, and maximizes restoration of gene expression change magnitudes under realistic temporal conditions, thereby enabling the velocities to accurately reflect biological reality. Alternatively, the transcriptomic velocities (*d***x**/*dĈ*)_*i*_ may be denoted by V_i_.

### Simulation of Single-Cell Data with Ground-Truth Velocities

We generated in silico single-cell RNA sequencing (scRNA-seq) datasets with known ground-truth transcriptomic velocities using scDesign3 [45]. This framework parametrically models gene expression as a time-dependent stochastic process, enabling analytical extraction of reference transcriptomic velocities.

Following the recommended practices in the official scDesign3 documentation and referencing the pancreatic endocrine differentiation dataset, we constructed a simulated dataset comprising 5,000 cells and 500 genes.

The core of scDesign3 employs Generalized Additive Models for Location, Scale, and Shape (GAMLSS) to model each gene’s expression distribution as a function of time. For each gene *s* ∈ {1, …, *G*}, the mean expression *μ*_*g*_(*t*) is parameterized as a smooth, non-linear function of time *t*. Although scDesign3 provides smooth mean functions *μ*_*g*_(*t*), it does not directly output transcriptomic velocity. We therefore analytically derived reference transcriptomic velocities through numerical differentiation of the fitted models. The instantaneous rate of change for gene *s* at time *t* is given by the temporal derivative:

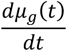

We approximated this derivative using a forward finite difference scheme with an infinitesimal perturbation *δ* = 10^−7^:

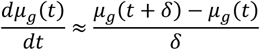

This procedure yielded simulated data with ground-truth transcriptomic velocities. To evaluate the impact of sampling density, we subsequently subsampled the simulated data using the following scheme:

Let 𝒞 = {*c*_1_, *c*_2_, …, *c*_*n*_} denote the set of *n* cells with time values 𝒯 = {*t*_1_, *t*_2_, …, *t*_*n*_}, where *t*_*i*_ ∈ [0,1] represents the time position of cell *c*_*i*_. For a specified number of sampling time points *m*, we defined a set 𝒫 of uniformly distributed sampling locations within the central time region:

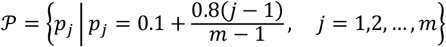

For each sampling point *p*_*j*_ ∈ 𝒫, we generated *K* = 500 target time values from a normal distribution centered at *p*_*j*_:

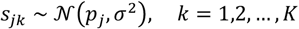

with bandwidth *σ* = 0.05 controlling the local selection range. Each target value *s*_*jk*_ selected the cell with the nearest time:

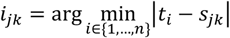

The initial index set ℐ_0_ = {*i*_*jk*_} potentially contained duplicates. The final sampled cell index set ℐ was obtained by retaining unique indices:

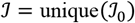

For the sampled dataset {*c*_*i*_: *i* ∈ ℐ}, we recorded the original sampling point *p*_*j*_ associated with each selected cell.

### Lineage Tree Node Sampling in *C. elegans*

Let 𝒞 denote the set of all cells with unambiguous sampling times. For each cell *c* ∈ 𝒞, let *t*(*c*) ∈ 𝒯 denote its time point, where 𝒯 = {*τ*_1_, *τ*_2_, …, *τ*_*i*_} is the set of all discrete time points. Let Λ = {*λ*_1_, *λ*_2_, …, *λ*_*k*_} be the set of distinct lineages present in 𝒞, where each *λ*_*i*_ represents a unique lineage node. For each lineage node λ ∈ Λ, define the cell set 𝒞_*λ*_. For each lineage node λ ∈ Λ, execute the following selection procedure.

Define the cell count function for lineage node *λ* at time point *τ*:

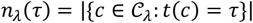

Find the maximum count across all time points:

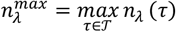

Identify the set of time points achieving this maximum:

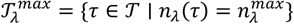

Define the candidate cells for lineage node λ as those belonging to the most frequent time point(s):

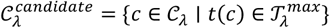

Select a single representative cell 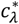 uniformly at random from the candidate set:

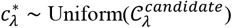

The final dataset 𝒞_*final*_ is the collection of all selected representative cells:

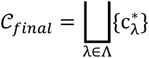

where ⨆ denotes the disjoint union, ensuring exactly one cell per lineage node. The resulting dataset contains |Λ| = 333 cells as stated. The selection method preserves the original temporal distribution by mode matching: for each lineage, it selects cells from the time point(s) with the highest sampling frequency.

### Quantification of Temporal Distribution Distance Using the Overlapping Coefficient

To quantify the temporal distance between tumor-associated macrophages (TAMs) and monocytes, we employed the overlapping coefficient (OVL), a non-parametric measure of similarity between two probability distributions. The OVL statistic is defined as the integrated area where two density distributions intersect, providing a robust metric for comparing temporal patterns across cell populations.

Let *X* = {*x*_1_, *x*_2_, …, *x*_*n*_} and *Y* = {*y*_1_, *y*_2_, …, *y*_*i*_} represent the sets of temporal measurements for TAMs and monocytes, respectively. The OVL was computed through the following procedure. First, to enable direct comparison, we constructed normalized density histograms for both distributions using identical binning parameters:

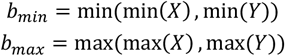

A uniform bin grid ℬ = {*b*_0_, *b*_1_, …, *b*_*k*_} was generated across [*b*_*min*_, *b*_*max*_] with *k* = 100 intervals of width Δ*b* = (*b*_*max*_ − *b*_*min*_)/*k*. Probability densities were estimated as:

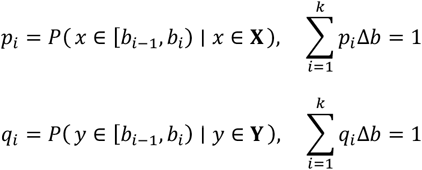

The overlapping coefficient was then calculated by integrating the minimum density across all bins:

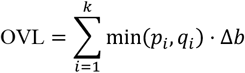

This yields a statistic bounded by [0,1], where 0 indicates disjoint distributions and 1 indicates identical distributions. The temporal distance metric was subsequently defined as 1 − OVL for visualization in Fig. 4.

### Quantifying Gene Contributions to Temporal Encoding

Let **X** ∈ ℝ^*n*×*t*^ denote the gene expression matrix for *n* cells and *m* genes, and **W** ∈ ℝ^*t*^ denote the vector of clock weights. Original time predictions **C** = **XW** are computed, and the isolated contribution of each gene is obtained by element-wise multiplication **G**_*ij*_ = **X**_*ij*_ ⋅ **W**_*j*_. Perturbed predictions **P**_*ij*_ = **C**_*i*_ − **G**_*ij*_, representing clock outputs with each gene’s effect removed, are then derived. Both **C** and **P** are globally min-max normalized to [0, 1], yielding:

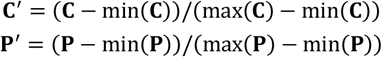

Two metrics quantify gene-specific impact across all cells (*i* = 1, …, *n*) for each gene (*j* = 1, …, *m*). Average contribution measures the overall effect and is calculated as the L2 norm:

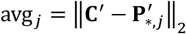

Peak contribution captures the maximum deviation in any single cell and is calculated as:

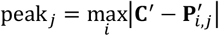

Both metrics are subsequently min-max normalized across genes to the [0, 1] range.

These contributions were evaluated across experimental groups: Fig. 4h depicts the average contribution per gene in both aTrem2 and IgG conditions, while Fig. 5h illustrates both average and peak contributions for individual genes.

### Gene-Specific Magnitude Scale of RNA Velocity

We computed a magnitude scale for each gene, quantifying the scaling factor required to align RNA velocities with gene-embedded time. Let ***V*** ∈ *R*^*n*×*g*^ denote the matrix of RNA velocities, and ***M*** ∈ *R*^*n*×*g*^ denote the corresponding matrix of transcriptomic velocities magnitudes inferred from CellDyc gene-embedded time (as described in equation (4)), where *n* is the number of cells and *s* is the number of genes.

For each gene *i*, the dynamic range of velocity activity was characterized by first computing the *p*-th percentile (*p* = 95.0 by default) of absolute velocity values across cells, denoted *τ*_*i*_. Velocities exceeding this threshold, 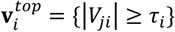, were selected to represent reliable velocity estimates, and their mean amplitude was calculated as 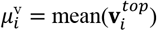. The corresponding baseline magnitude of transcriptomic velocities was quantified by the per-gene mean 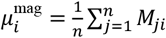. The gene-specific magnitude scale factor *s*_*i*_ was then derived as the ratio of baseline magnitude to velocity amplitude:

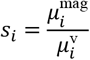

This procedure yielded the magnitude scale vector ***S*** = (*s*_1_, *s*_2_, …, *s*_*g*_).

### Application of Other Methods

#### Waddington-OT and Moscot

Both Waddington-OT [39] and Moscot [40] compute cell-to-cell transition matrices. These matrices were projected onto the embedding space using CellRank’s RealTimeKernel [44].

#### CellRank

We performed downstream fate probability analysis for all methods using CellRank [43, 44]. Transition matrices from Waddington-OT [39] and Moscot [40] were analyzed using RealTimeKernel [44], whereas those from scVelo [27] and cellDancer [28] were processed with VelocityKernel [44].

#### scVelo and cellDancer

Velocity calculations for scVelo [27] and cellDancer [28] were carried out in accordance with the recommended procedures outlined in their respective official documentations.

### scRNA-seq Data and Pre-processing

All single-cell RNA sequencing (scRNA-seq) datasets utilized in this study were obtained from publicly available sources. Data pre-processing includes filtering, normalization, selection of highly variable genes, principal component analysis (PCA), and construction of a neighborhood graph. Following pre-processing, the top 2,000 highly variable genes and the neighborhood graph were retained for subsequent analysis.

#### *C. elegans* embryonic data

The raw dataset was originally published by Packer et al [8]. As described in the “Lineage Tree Node Sampling in *C. elegans*” section, we selected 333 cells.

#### Zebrafish embryogenesis data

The raw dataset was generated by Farrell et al [9]. We adopted the cell selection method of Lange et al. [43], obtaining 2,341 cells.

#### Glioblastoma-associated monocyte differentiation data

The raw dataset was initially described by Kirschenbaum et al [42]. We followed the alignment procedure of Proks et al. for MARS-seq data to derive spliced and unspliced reads for RNA velocity comparisons [68]. Cell selection was performed according to Kirschenbaum et al. to isolate monocyte differentiation-related populations, yielding 3,108 cells.

#### Mouse gastrulation erythropoiesis data

The raw dataset was published by Pijuan-Sala et al [10]. We followed the approach of Li et al. [28], selecting cells from embryonic stages E6.75 to E8.5 based on annotated cell type classifications: haematoendothelial progenitors, blood progenitors 1/2, and erythroid 1/2/3. Cells with pronounced outlier characteristics in low-dimensional space were excluded, resulting in a final dataset of 12,324 cells.

#### Mouse embryonic fibroblast reprogramming data

The raw dataset was reported by Biddy et al [11]. We employed the subset curated by Lange et al [43]. (48,515 cells) and subsequently filtered to retain only cells with definitive lineage markers indicative of either successful reprogramming or dead-end trajectories, resulting in 3,049 cells.

## Extended Data Figures

**Extended Data Fig. 1.**
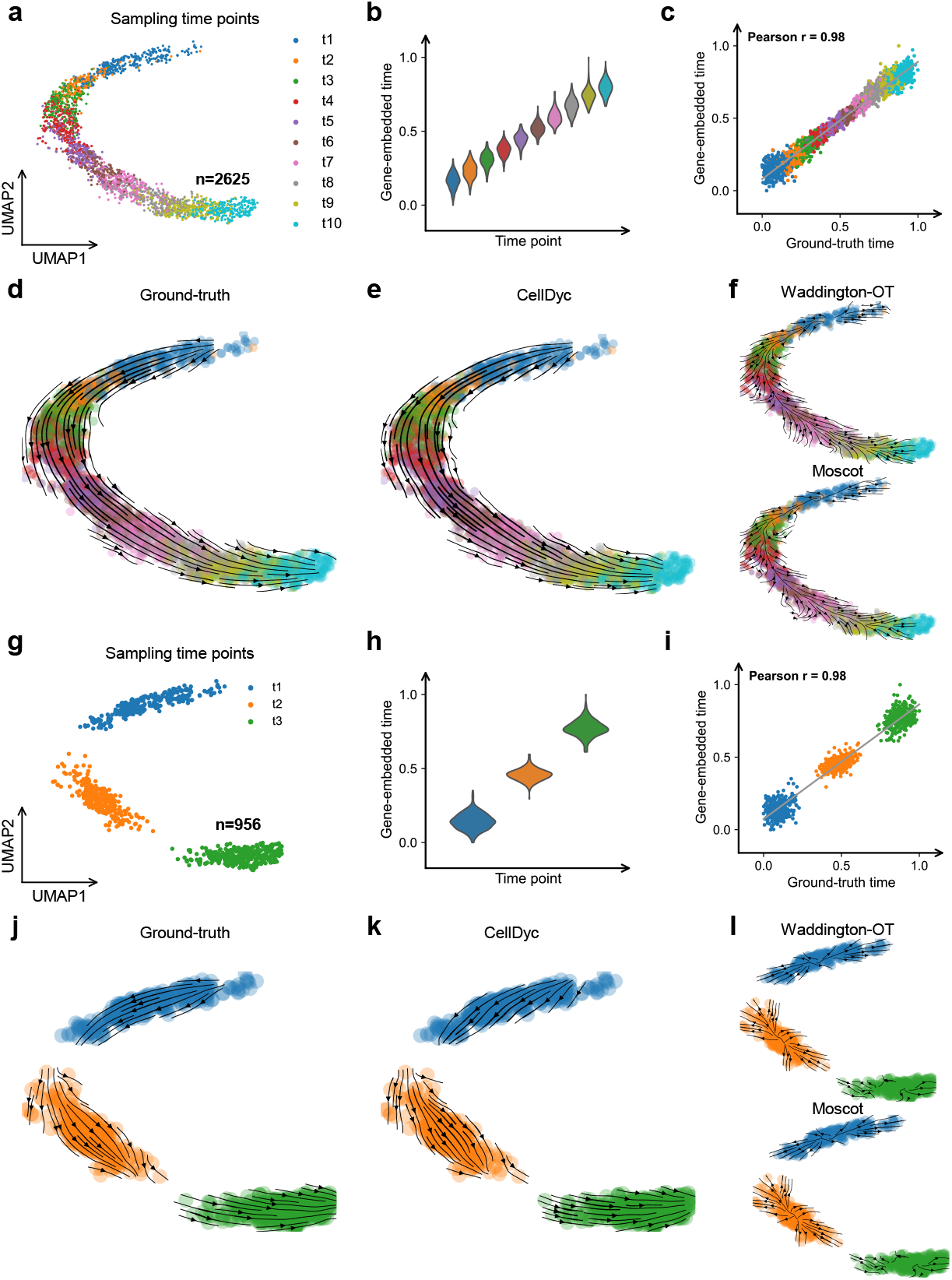
Performance benchmarking of CellDyc in simulations with varying temporal sampling densities (Related to Fig. 2). **a**, Temporal sampling at ten time points according to a normal distribution, resulting in 2,625 cells. **b**, Violin plots of predicted gene-embedded time across sampling time points. **c**, The correlation between predicted gene-embedded time and the ground-truth time, Pearson’s correlation coefficient is indicated. **d,e**, Projection of transcriptomic velocities onto UMAP. (**d**) Ground-truth velocities. (**e**) CellDyc-predicted velocities. **f**, Projection of cell state transitions onto UMAP: (top) Waddington-OT, (bottom) Moscot. **g**, Temporal sampling at three points according to a normal distribution, resulting in 956 cells. **h**, Violin plots of predicted gene-embedded time across sampling time points. **i**, The correlation between predicted gene-embedded time and the ground-truth time, Pearson’s correlation coefficient is indicated. **j,k**, Projection of transcriptomic velocities onto UMAP. (**j**) Ground-truth velocities. (**k**) CellDyc-predicted velocities. **l**, Projection of cell state transitions onto UMAP: (top) Waddington-OT, (bottom) Moscot.

**Extended Data Fig. 2.**
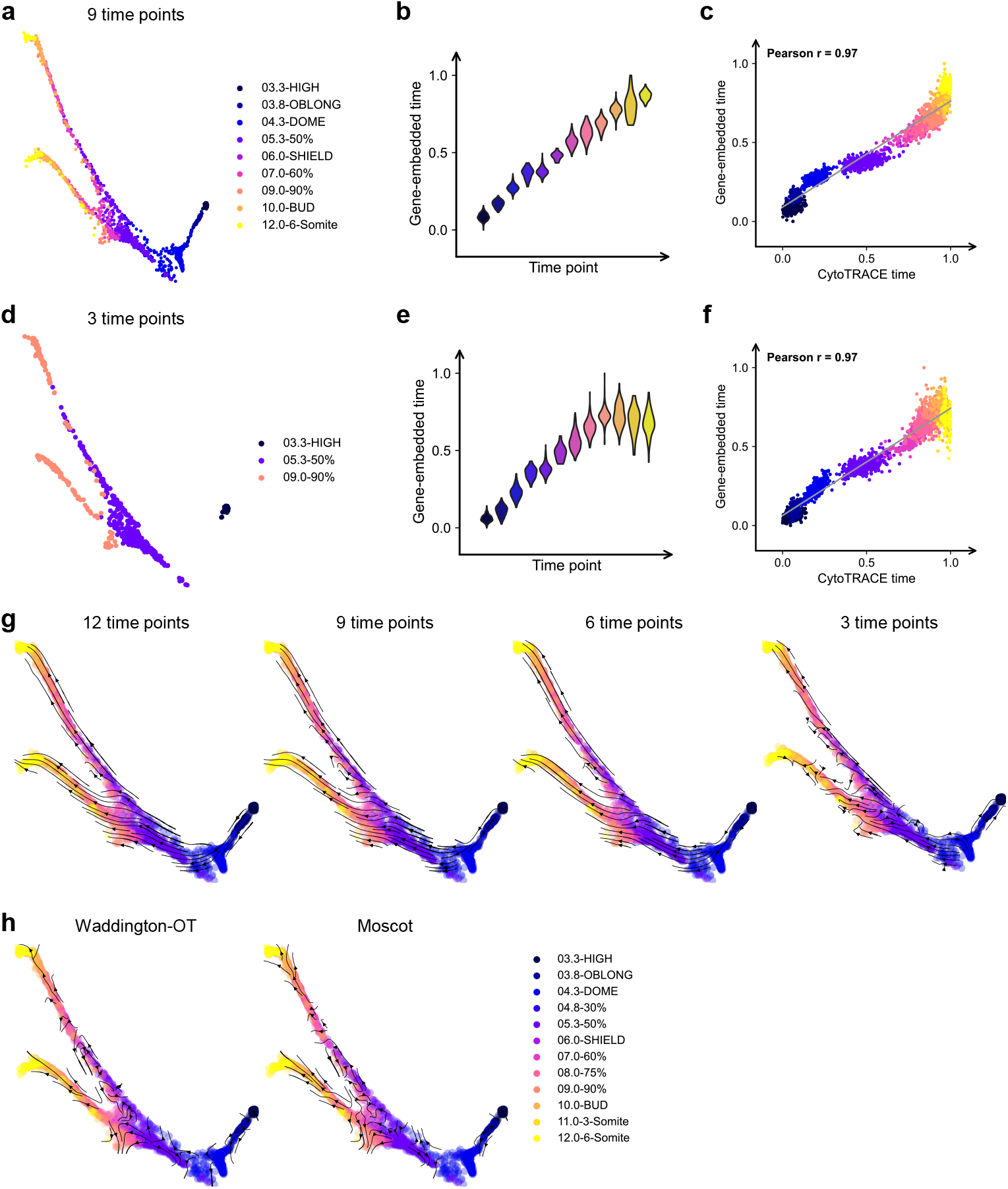
Reconstruction of zebrafish developmental dynamics and masked time points under variable sampling conditions (Related to Fig. 3). **a**, Nine time points selected for CellDyc training are shown. **b**, Violin plots of predicted gene-embedded time across sampling time points from the selected 9-time-point model. **c**, The correlation between 9-time-point model predicted gene-embedded time and CytoTRACE time. Pearson’s correlation coefficient is indicated. **d**, Three time points selected for CellDyc training are shown. **e**, Violin plots of predicted gene-embedded time across sampling time points from the selected 3-time-point model. **f**, The correlation between 3-time-point model predicted gene-embedded time and CytoTRACE time. Pearson’s correlation coefficient is indicated. **g**, Projection of transcriptomic velocities derived from different models onto Force-directed layout. From left to right: 12-time-point model, 9-time-point model, 6-time-point model, and 3-time-point model. **h**, Projection of cell state transitions onto Force-directed layout: (left) Waddington-OT, (right) Moscot.

**Extended Data Fig. 3.**
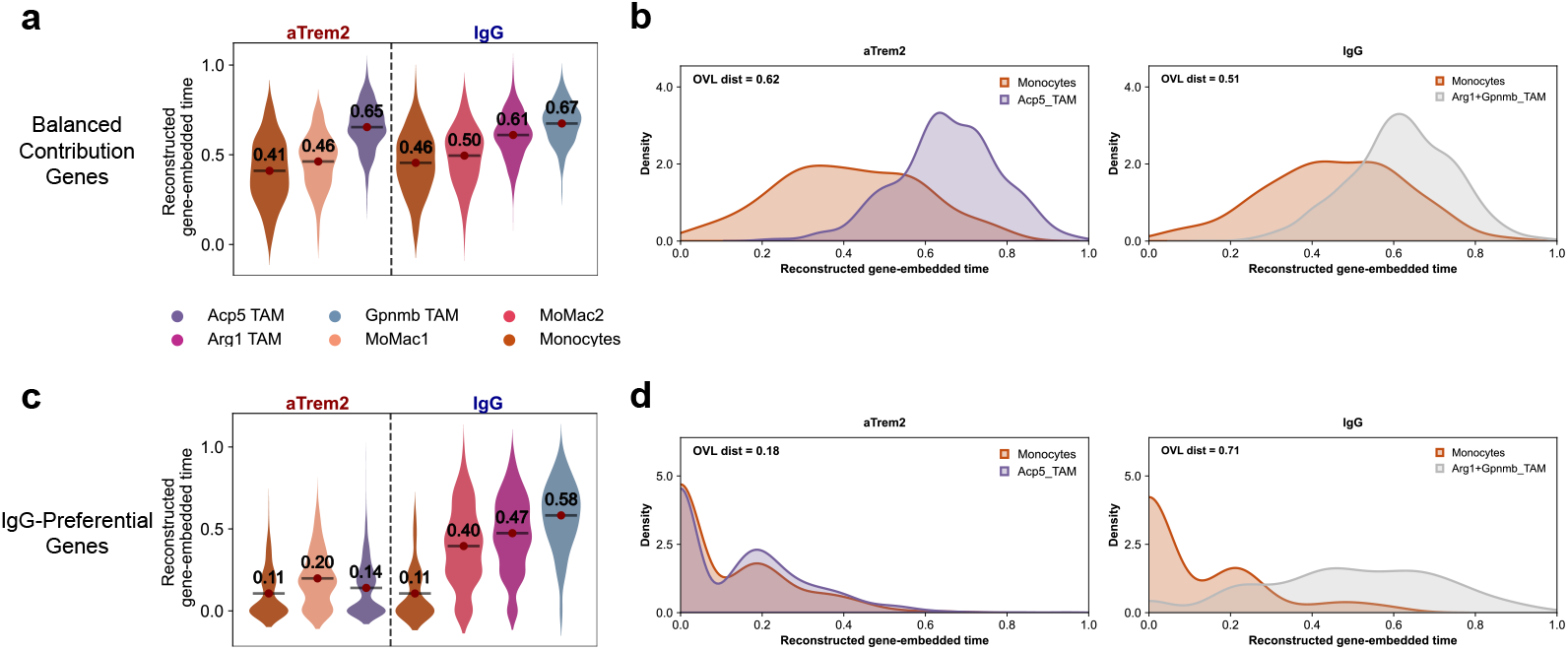
Reconstruction of gene-embedded time using identified “Balanced Contribution” and “IgG-Preferential” gene sets (Related to Fig. 4). **a**, Violin plots of gene-embedded time reconstructed using the nine “Balanced Contribution Genes” across cell types under aTREM2 treatment versus IgG control conditions. **b**, Distributions of the reconstructed gene-embedded time shown in (**a**) for monocytes and TAMs under aTREM2 treatment versus IgG control conditions, with OVL distance between TAMs and monocytes indicated. Left: aTREM2 treatment; Right: IgG control. **c**, Violin plots of gene-embedded time reconstructed using the four “IgG-Preferential Genes” across cell types under aTREM2 treatment versus IgG control conditions. **d**, Distributions of the reconstructed gene-embedded time shown in (**c**) for monocytes and TAMs under aTREM2 treatment versus IgG control conditions, with OVL distance between TAMs and monocytes indicated. Left: aTREM2 treatment; Right: IgG control.

**Extended Data Fig. 4.**
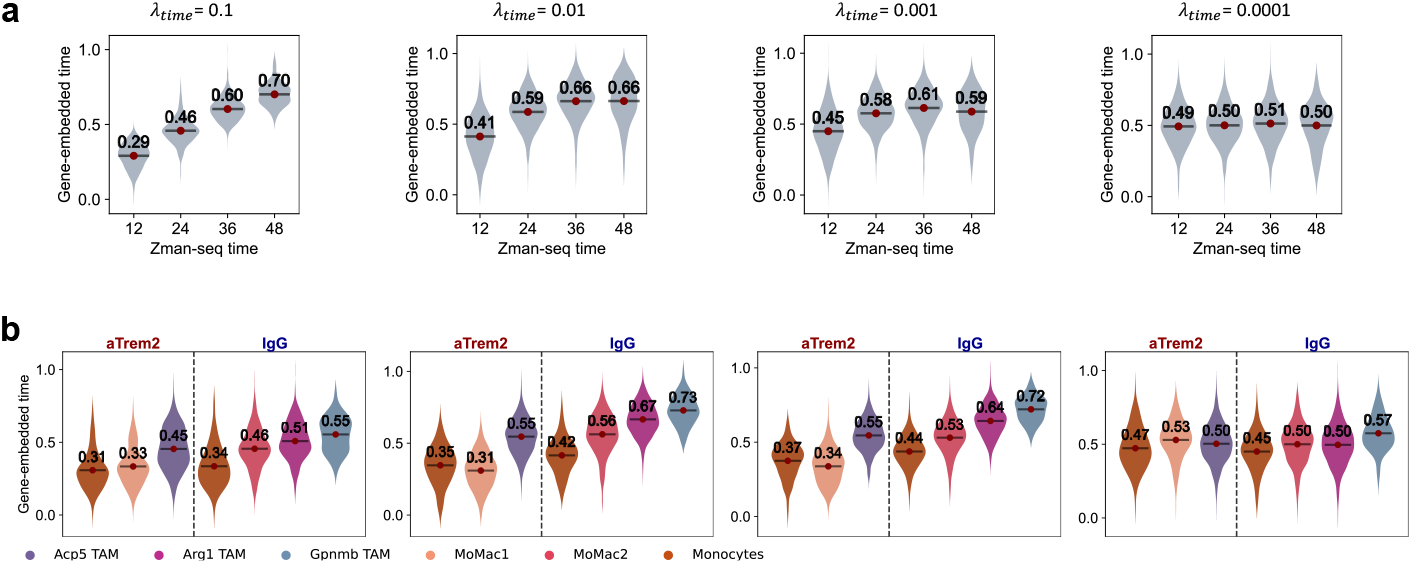
Sensitivity analysis of the time-loss weight parameter *λ*_*time*_ and its impact on gene-embedded time prediction (Related to Fig. 4). **a**, Violin plots of the predicted gene-embedded time with different *λ*_*tttt*_ settings across Zman-seq time points. The plots correspond to *λ*_*tttt*_ settings of 0.1, 0.01, 0.001, and 0.0001 from left to right. **b**, Violin plots.

**Extended Data Fig. 5.**
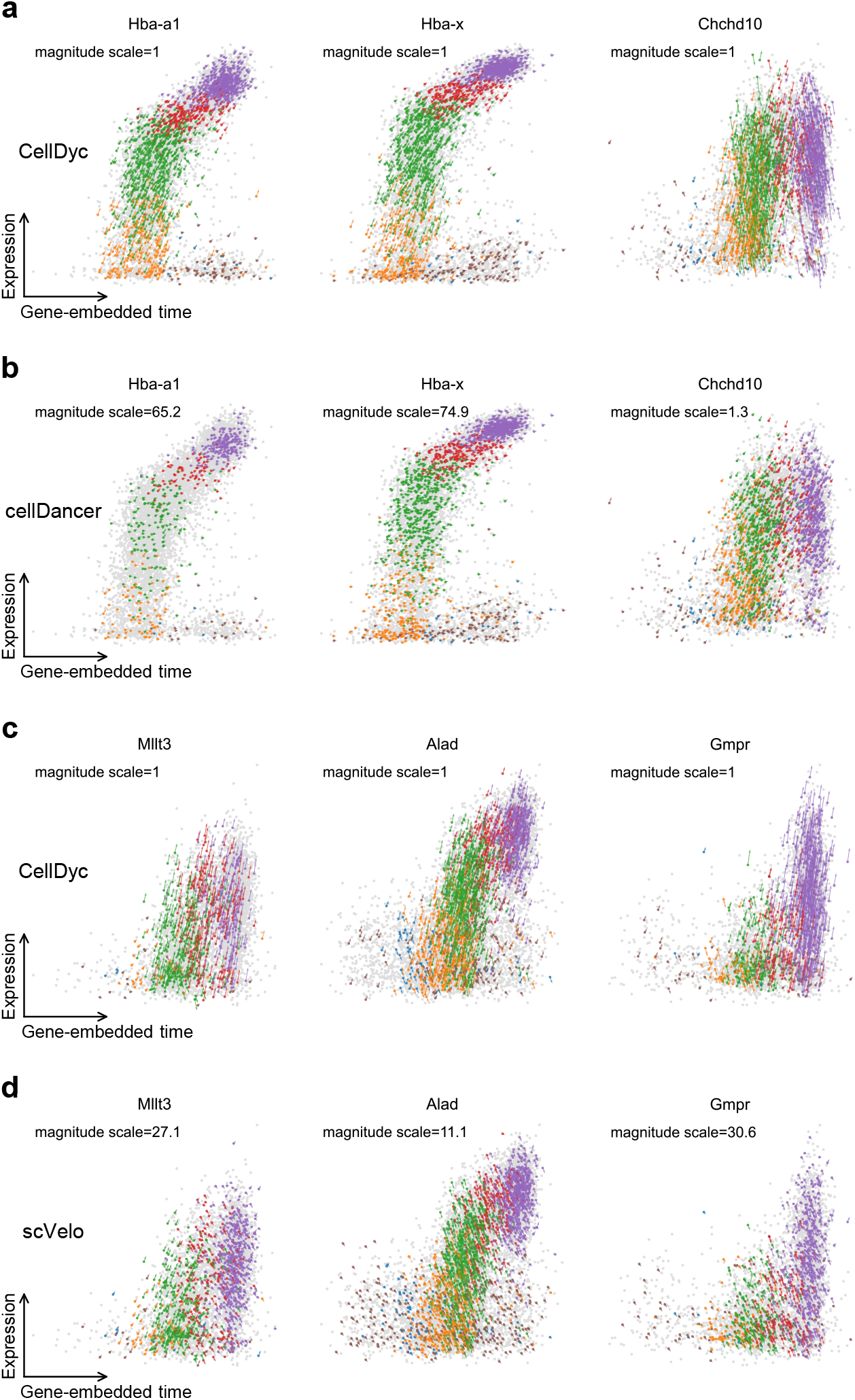
Comparative analysis of single-gene velocity dynamics inferred by CellDyc, cellDancer, and scVelo (Related to Fig. 5). **a,b**, Comparison of velocity dynamics for selected genes between CellDyc and cellDancer. (**a**) CellDyc. (**b**) cellDancer. **c,d**, Comparison of velocity dynamics for selected genes between CellDyc and scVelo. (**c**) CellDyc. (**d**) scVelo.

**Extended Data Fig. 6.**
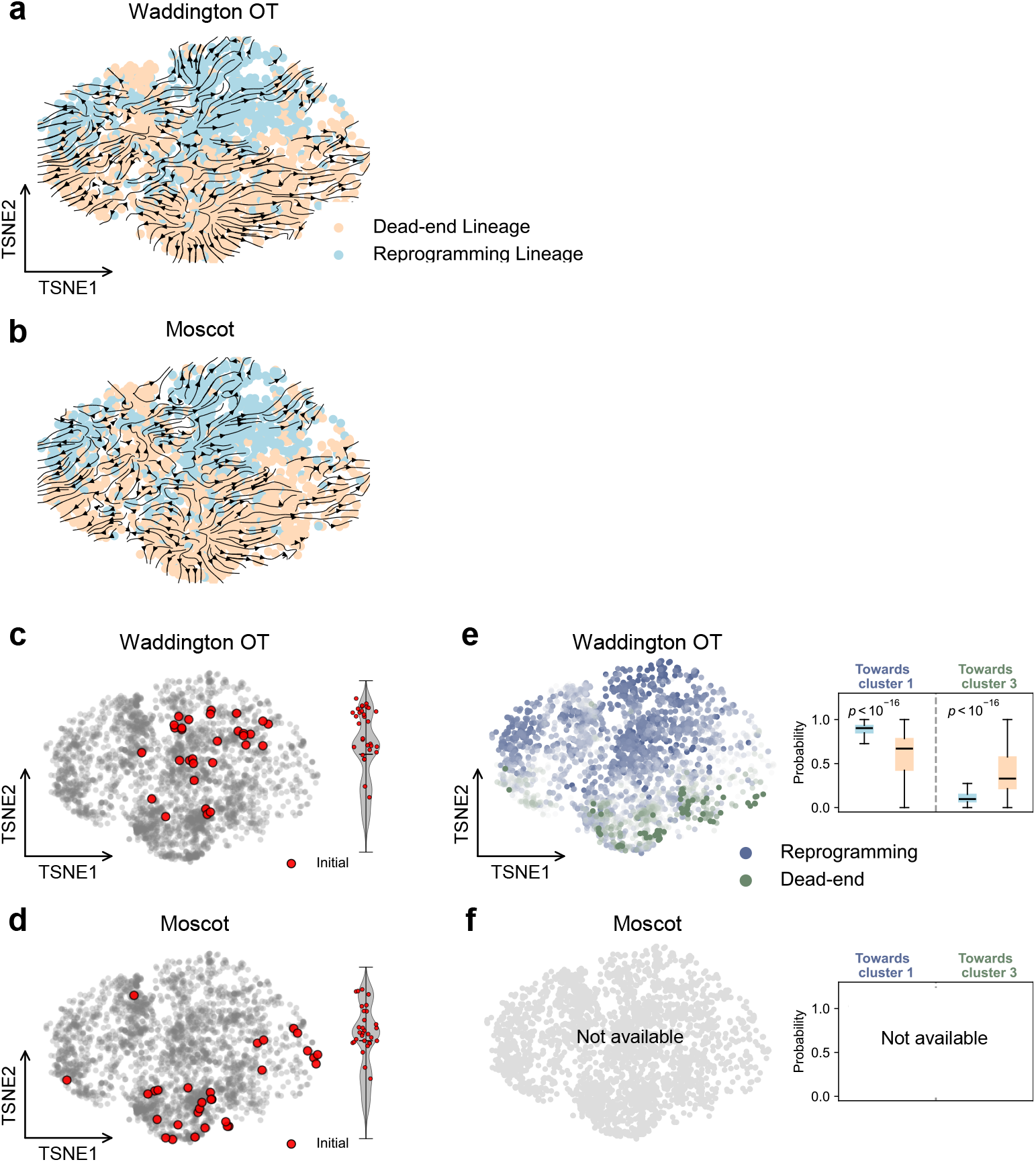
Cell fate probability inference in fibroblast reprogramming using Waddington-OT and Moscot-derived transitions (Related to Fig. 6). **a,b**, Projections of cell state transitions derived from Waddington-OT (**a**) and Moscot (**b**) onto the t-SNE. Cells are colored by lineage barcodes from CellTagging lineage tracing. **c,d**, Initial (red) and terminal (blue) macrostates determined by CellRank based on the cell state transitions from Waddington-OT (**c**) and Moscot (**d**), as in Fig. 6e. **e,f**, Cell fate probabilities for cluster 1 versus cluster 3, computed using the state transitions from Waddington-OT (**e**) and Moscot (**f**) with manually anchored terminal states, as in Fig. 6f. Left: t-SNE projection showing the probability of each cell reaching the combined macrostates of cluster 1 and cluster 3. Right: Boxplots of fate probabilities stratified by terminal states.

## Notes

### Competing Interest Statement

The authors have declared no competing interest.

